# Spatial analysis of Intraductal Papillary Mucinous Neoplasms reveals secretory cell-enriched neighborhoods

**DOI:** 10.64898/2026.06.16.732658

**Authors:** Amelia T. Cephas, Brenda Jarvis, Katherine Gell, Christopher P. Taranto, Maelle Batardiere, Sasha Sapon-Cousineau, E. Danielle Dean, Aatur D. Singhi, Marcus C.B. Tan, Vincent Q. Trinh, Kathleen E. DelGiorno

## Abstract

Pancreatic ductal adenocarcinoma (PDAC) is currently the third leading cause of cancer-related deaths in the United States. Intraductal papillary mucinous neoplasms (IPMNs) are neoplastic lesions of ductal origin that seed 10-25% of PDAC. There are currently no markers that distinguish between IPMN that will remain benign and those that will progress to cancer. A heterogenous population of secretory cells, including chemosensory tuft cells and hormone-expressing enteroendocrine cells (EECs), form during metaplasia and neoplastic progression in the pancreas, but the relevance of these populations as it relates to IPMN progression is not well characterized. Here, we performed spatial transcriptomics as well as multiplex immunostaining and spatial statistics on surgically resected IPMN from 60 patients to characterize these populations in all subtypes (gastric foveolar, intestinal, pancreatobiliary) and grades (low-grade, high-grade, invasive). We found that POU2F3^+^ tuft-like cells, CHGA^+^ EECs, and a subset of pancreatic endocrine cells (ɑ and γ cells) were present in all types of IPMN. Further, serotonin-expressing enterochromaffin cells made up the bulk of EECs in low-grade disease. Enterochromaffin, tuft-like, and glucagon-expressing alpha cells were not evenly distributed and instead were significantly enriched in a spatial manner, which is overlooked using conventional whole tissue quantification approaches. Tuft-like cell clusters were enriched with monocytes and resident memory T cells and anti-correlated to activated fibroblasts (myCAFs, iCAFs). Overall, these secretory cell clusters may reflect clonal expansion resulting in formation of distinct stromal niches with unknown consequences for disease progression.

## INTRODUCTION

Pancreatic ductal adenocarcinoma (PDAC) is an aggressive and lethal disease with a 5-year survival rate of only 13%[1]. It is predicted to become the second leading cause of cancer related deaths in the United States by the year 2040[2]. Further, the incidence of PDAC is rising in young adults aged 15-39 years old, influenced in part by rising obesity incidence within this age group[3,4]. With its complex and aggressive tumor biology and the fact that patients often present as asymptomatic, PDAC is very difficult to detect, and thus is diagnosed at late stages[1]. From histological and genetic studies, most PDACs arise from microscopic acinar to ductal metaplasia (ADM) lesions that progress to pancreatic intraepithelial neoplasia (PanIN)[5–8]. A subset of PDAC, however, arise from radiologically detectable, often indolent, macroscopic lesions called intraductal papillary mucinous neoplasia (IPMN)[8,9].

IPMN can be classified by histological subtype (variant), which includes intestinal, gastric foveolar (GF), and pancreatobiliary (PB). Intestinal IPMN resemble intestinal villi with tall columnar epithelial cells, whereas GF-IPMN resemble gastric foveolae. PB-IPMN exhibit cholangiopapillary-like architecture with thin, branching papillae. GF-IPMNs are predominately found in the branch ducts and are low-grade, characterized by mild cellular atypia with generally preserved ductal structure and require surveillance[8,10]. In contrast, intestinal and PB-IPMN are found in both branched and main ducts and are generally higher-grade lesions. Although up to 90% of IPMNs are detected by routine imaging, curative-intent surgery is the only therapeutic option [11,12]. Pancreatic surgery, however, carries substantial risk of major complications and mortality. Because there are no available tests to predict the grade of dysplasia preoperatively to estimate risk of progression, management of high-risk cysts relies heavily on international consensus guidelines and a multidisciplinary approach[13].While risk stratification strategies offer insight into the radiographic, clinical, and biochemical features of IPMN, they do not accurately predict disease progression, making the study of cellular changes in IPMN progression critical to develop better diagnostics[14,15].

We have previously shown that several cell types emerge in the pancreas during ADM[16], an early event in tumorigenesis whereby acinar cells, the digestive enzyme-producing cells of the exocrine pancreas, transdifferentiate into a heterogeneous population of ductal cells. For example, hormone-producing enteroendocrine cells (EECs) and chemosensory tuft cells are rare in the normal pancreatic ductal epithelium; however, we have shown that they form *de novo* in ADM[16–18]. EECs are a subset of endocrine cells that play vital roles in nutrient absorption and motility in the gastrointestinal tract. EECs are abundant in ADM but decrease in number with neoplastic progression, consistent with loss of epithelial differentiation with disease progression[17]. Although pathological studies have described endocrine cells—specifically those populations constituting pancreatic islets—as associated with IPMN[19], they are understudied in this disease context.

Like EECs, tuft cells expand in injury or oncogene-induced ADM, decrease in abundance with neoplastic progression, and are absent from invasive disease[20]. Consistent with a differentiated cell type, we have shown that these chemosensory cells play a functional role in pancreatic tumorigenesis, mitigating inflammation through secretion of factors like PGD_2_ [21,22], inhibiting tumor progression. Whether the presence of EECs and tuft cells distinguish IPMN subtype, serve as an indicator of tumor progression, or function in disease progression, is unknown. This study employed spatial transcriptomics, multiplex immunostaining, and spatial statistics to investigate tuft, EEC, and islet endocrine cell abundance in patient IPMN. Here, we show that these secretory cell populations expand in these cysts. Specifically, tuft-like cells, serotonin-expressing enterochromaffin cells, and glucagon-expressing ɑ cells regionally expand, forming clusters and distinct stromal neighborhoods, revealing a new layer of previously unappreciated epithelial heterogeneity and niche specification.

## MATERIALS AND METHODS

### Study Cohort

A cohort of 60 patients with intraductal papillary mucinous neoplasms (IPMNs) was selected from the Vanderbilt University Medical Center institutional cohort under institutional review board approval (IRB #110061, **Table S1**). Specimens analyzed spanned the spectrum of disease progression, including low-grade dysplasia (LG; n = 38), high-grade dysplasia (HG; n = 22), and invasive carcinoma (Inv; n=14). Malignant or invasive carcinoma was documented in 14 of 60 cases (23.3%), including 4 low-grade and 10 high-grade IPMN cases. Histologic subtypes represented included intestinal (n = 9), gastric foveolar (GF; n = 23), pancreatobiliary (PB; n = 13) and mixed (n=2) IPMN. Histologic subtype was unspecified in 13 cases, six of which were malignant.

## RESULTS

### IPMN express Tuft cell and EEC markers

Tuft cells and EECs are found in ADM, and their abundance is negatively correlated with PanIN progression; however, it is not known whether these populations distinguish IPMN subtype or are indicators of IPMN progression. To identify secretory cell transcriptional programs in IPMN, we first performed NanoString GeoMx spatial transcriptomic profiling (**Figure 1A**). We analyzed four patient samples—two intestinal-type and two gastric-foveolar (GF)-IPMN. Regions of interest (ROIs; n=95 or 23-24/sample) were defined using a cell morphology marker (γ-actin) and a single marker to delineate regions containing secretory cells (synaptophysin, SYP). Using principal component analysis (PCA), we found that all ROIs (∼513-516 μm) clustered by patient sample with both GF-IPMN samples clustering more closely than with intestinal type IPMN (**Figure 1B**). ROIs with islet and acinar tissue, as determined by cellular morphology, expressed expected markers, INS/GCG, and CPA1/PRSS2, respectively, validating our analysis (**Figure 1C**). The top 50 most variable genes across patients were commensurate with molecular subtype as determined by previously published markers[23] and pathologist evaluation (VQT). For instance, GF samples expressed *CLU, GKN1/2,* and *MUC6*[23]. Genes unique to intestinal samples included *CLDN3/4*[24], *CXCL5*[25], and *TGM2*[26] (**Figure 1D**), which are involved in fibrosis and inflammation. One HG intestinal sample (IPMN-Intestinal 1) showed high expression of the tuft cell marker *POU2F3* and a LG gastric (IPMN-Gastric 1) and HG intestinal (IPMN-Intestinal 1) sample demonstrated high expression of broad endocrine marker *CHGA* at both the gene and protein levels (**Figure 1E-F**). Given that *POU2F3* expression alone does not establish tuft cell identity, we refer to these cells as POU2F3^+^/tuft-like throughout the study.

**Figure 1.**
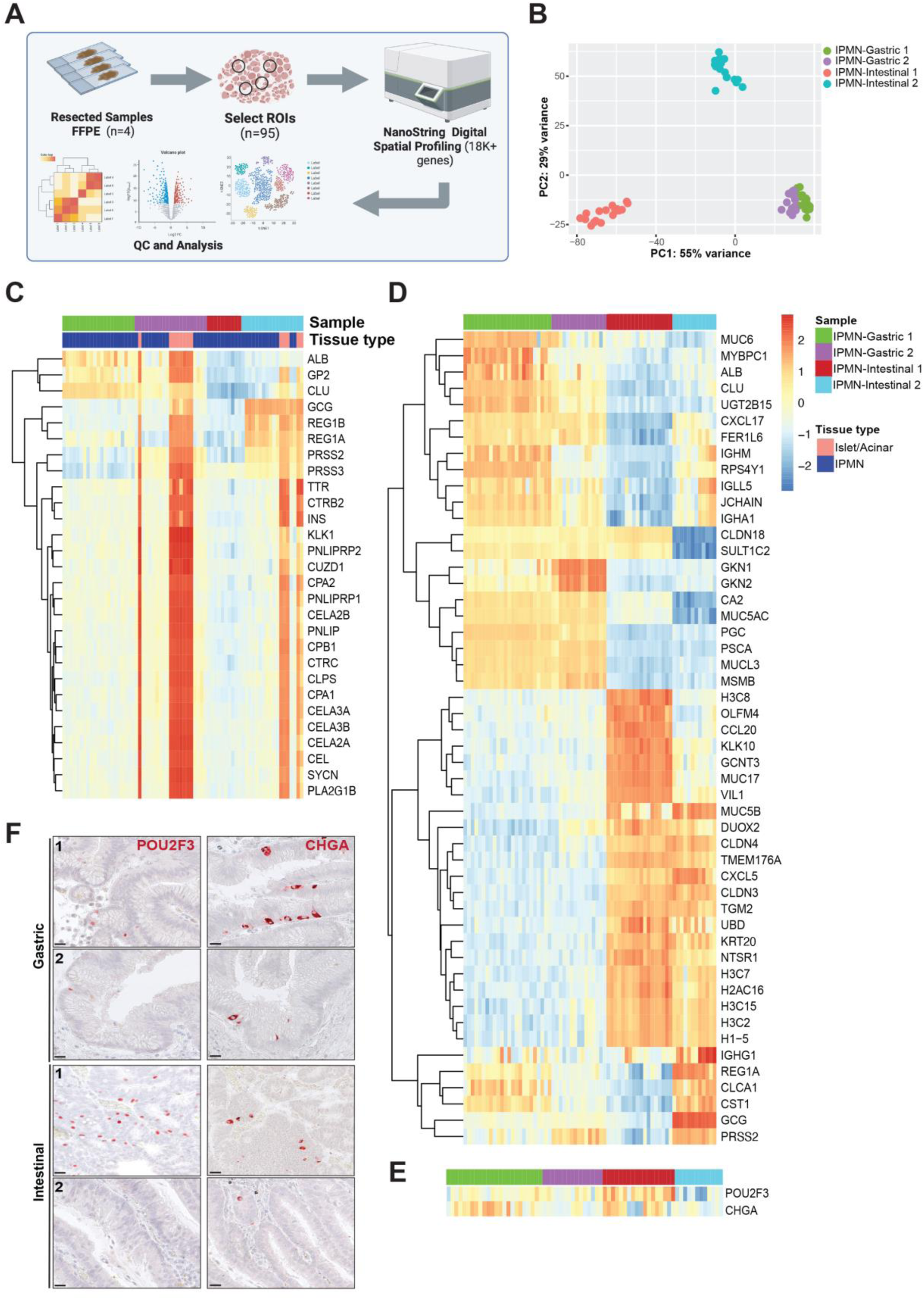
Spatial transcriptomics reveal tuft and EEC markers in IPMN. (A) Schematic of methods for NanoString GeoMx Whole Transcriptome Atlas (WTA) analysis. (B) Principal component analysis (PCA) plot of regions of interest (ROIs) per patient sample; n=4 patients and n=95 ROIs. Representative regions: Gastric 1 (LG), Gastric 2 (LG/HG), Intestinal 1 (HG/INV), and Intestinal 2 (HG). (C) Unsupervised heatmap of gene expression within ROIs across IPMN samples (n=93 ROIs). (D) Unsupervised heatmap showing the top 50 most variable genes across IPMN ROIs, islet/acinar encompassing ROIs excluded (n=80 ROIs). (E) Expression of tuft (*POU2F3*) and EEC (*CHGA*)-related transcripts. (F) Immunohistochemistry (IHC) staining of POU2F3 and CHGA in IPMN patient samples used for NanoString analysis. Scale bars, 50 μm. FFPE, formalin-fixed, paraffin-embedded; QC, quality control.

### Tuft-like cells are focally enriched in IPMN

To determine the abundance of tuft-like and endocrine/EECs in IPMN and whether prevalence correlates with molecular subtype or disease progression, we next performed multiplex immunohistochemistry (MxIHC) and quantitative statistical analyses for tuft cell marker POU2F3 and broad endocrine marker CHGA across 36 and 38 IPMN samples from the VUMC cohort, respectively (**Figure 2**). For POU2F3, we expanded the cohort by performing standard immunohistochemistry (IHC) on 8 additional samples. In total, 44 samples were analyzed, representing 42 unique patients, including two matched samples. The IPMN analyzed represented the full spectrum of disease progression including LG (n=18), HG (n=19), and invasive (INV, n=7) tumors. These samples included intestinal (n=6), GF (n=21), and PB (n=17) IPMN (**Table S1**). POU2F3⁺ IPMN cells were abundant in both LG and HG IPMN in comparison to INV disease, and in GF IPMN as compared to intestinal and PB subtypes, though these differences did not reach significance (**Figure S1A**). However, pathologist review of immunostains (by VQT) determined that POU2F3^+^ cells were not evenly distributed and focally aggregated in all subtypes of IPMN (**Figure 3A)**. To quantitatively test this observation, we performed spatial statistics using Ripley’s K[27–29]. Ripley’s K is a spatial statistical analysis that tests whether such patterns are nonrandom by comparing the observed distribution of positive cells (obs) to a random distribution (theo) across the same tissue. Ripley’s K was transformed to Ripley’s L for interpretation, where L(obs-theo) values above 0 indicate clustering, and maxL represents the strongest clustering strength observed across the distances tested. Thirty-two samples contained sufficient POU2F3^+^ cells for analysis and violin plots of L(obs–theo) showed values above 0 in nearly all samples, indicating clustering across the cohort (**Figure 3B)**. Further, 14/32 samples showed significant spatial enrichment by maxL (p <0.05) (**Figure S2A**). To corroborate spatial findings, we performed an additional statistical analysis, Moran’s I[30], to determine whether POU2F3⁺ IPMN cells were spatially autocorrelated, meaning whether nearby cells showed similar POU2F3 expression patterns[31]. Among the samples that met the criteria for analysis, 16/27 exhibited significant positive spatial autocorrelation, with Moran’s I values >0.1 (p<0.05). Moran’s I values ranged from <-0.20 to 0.33, and strong positive spatial autocorrelation (>0.3) was observed in three samples (**Figure S2B-D**); eight samples showed significant clustering by both maxL and Moran’s I (**Figure S2E**).

**Figure 2.**
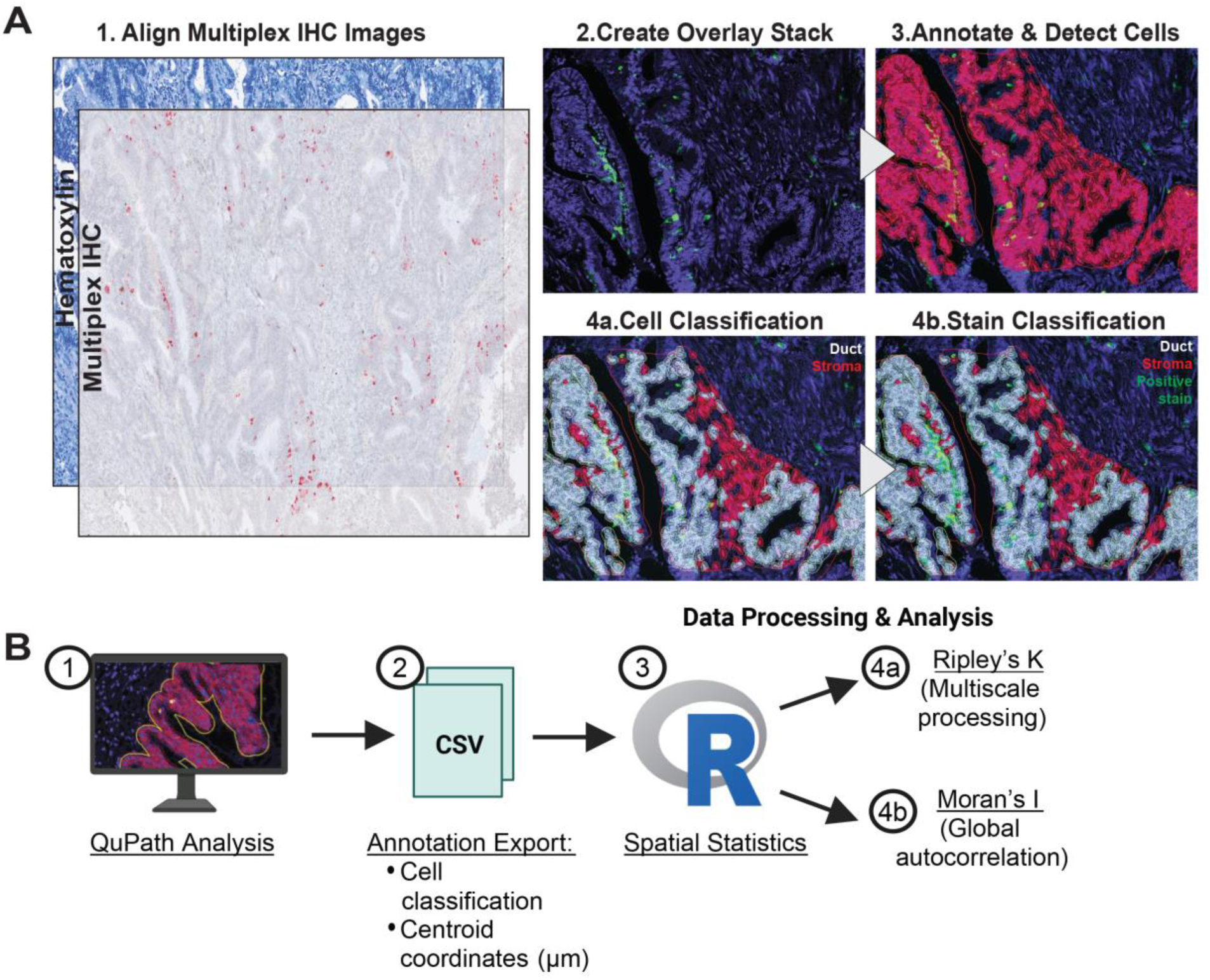
Cell quantification and spatial statistics methods. (A) 1-Multiplex-immunohistochemistry (mxIHC) staining was performed using hematoxylin counterstain and AEC chromogen detection to visualize POU2F3 and CHGA expression. 2- Hematoxylin and mxIHC images were aligned and overlayed with QuPath. 3-Ducts were manually annotated. 4a-Cells were detected using the Cell Detection function. 4b-Object classifiers for each protein were defined and applied. Immunofluorescence and DAB-stained tissues followed the same quantification pipeline, omitting the overlaying step in QuPath. (B) Steps for spatial statistics analysis using Ripley’s K and Moran’s I in R programming language.

**Figure 3.**
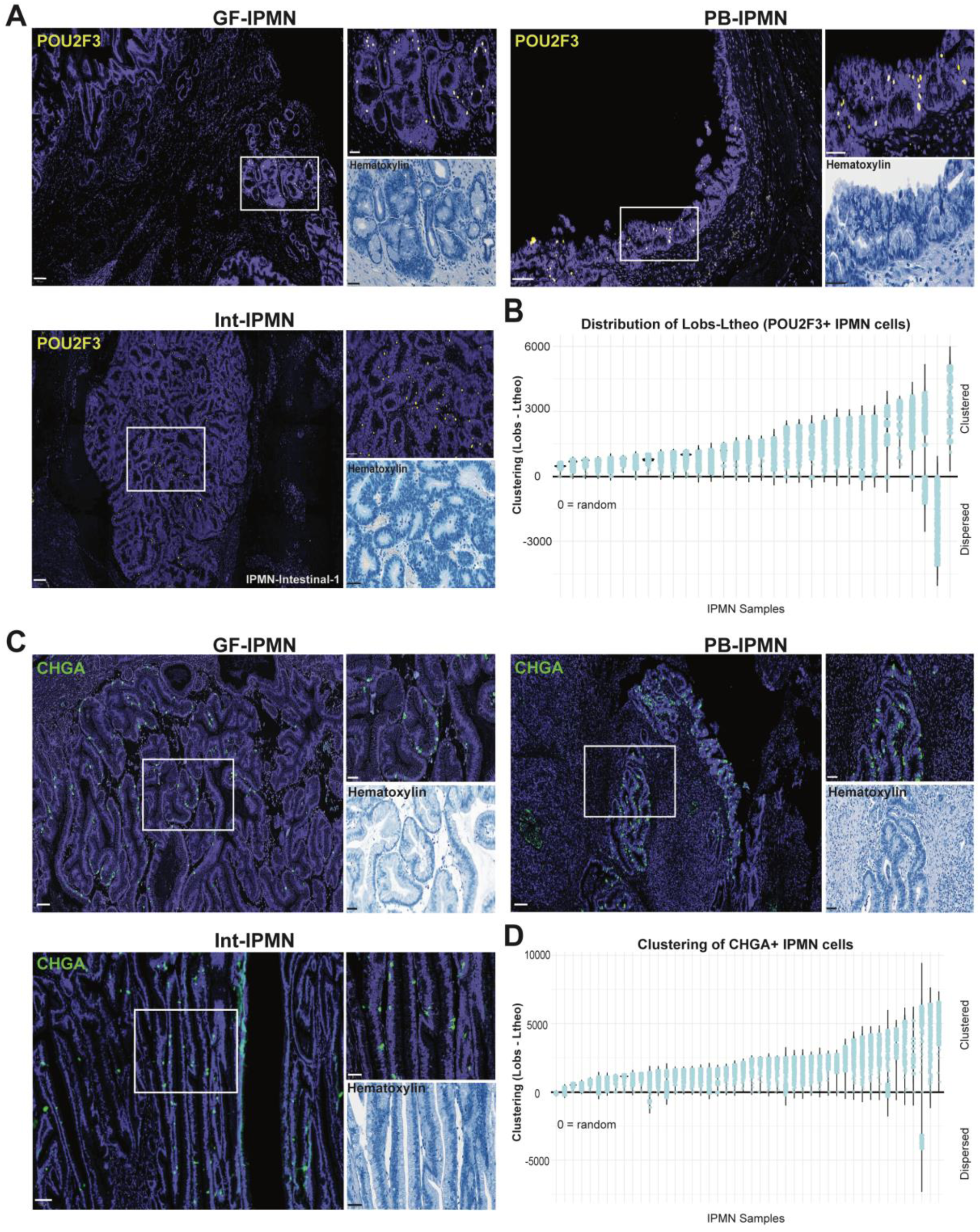
Tuft-like and endocrine cells are focally enriched in IPMN. (A) Representative pseudo colored IHC and hematoxylin images showing focal enrichment of POU2F3^+^ nuclei (yellow) in gastric foveolar (GF), pancreatobiliary (PB) and intestinal (Int) IPMN. (B) Violin plot with the distribution of L(obs-theo) of POU2F3^+^ nuclei across tested spatial distances in each IPMN sample (n=32). The black line marks 0, where observed clustering equals random expectation; values above or below 0 indicate clustering or dispersion, respectively. Samples are ordered by increasing clustering strength. (C) Representative pseudo colored IHC and hematoxylin images showing focal enrichment of CHGA^+^ cells (green) for GF, PB and Int IPMN. (D) Violin plot with the distribution of L(obs-theo) of CHGA^+^ cells across tested spatial distances in each IPMN sample (n=46). Scale bars, 100 μm; insets, 50 μm.

To examine POU2F3 expression in a second cohort of patient samples, we performed immunostaining on tissue microarrays generated at the University of Pittsburgh (TMAs, n=3 arrays; 236 patient samples). This cohort included normal pancreas, and multiple IPMN subtypes (PB, GF, GF-PB, and Int) and grades (LG, HG, and INV). After excluding non-evaluable samples, normal ducts and ADM, 131 samples were included in the final analyses (PB, n=27; GF, n=68; GF-PB, n=13; Int, n=9) and grades (LG, n=71; HG, n=46) as well as PDAC (INV, n=14). No significant differences in the abundance of percent POU2F3^+^ nuclei were observed across IPMN grade or subtype (**Figure S3A**). TMAs were then scored by two blinded pathologists (VQT, SSC) based on POU2F3^+^ staining of IPMN as absent (0), isolated cells (1), or small clusters (2). The percent of POU2F3^+^ nuclei significantly correlated with TMA score, such that samples with more abundant staining exhibited more POU2F3^+^ nuclei clustering (**Figure S3B**). POU2F3^+^ TMA scores were significantly lower in INV samples by grade (p<0.05) and trended towards lower in Int samples by variant. This suggests that POU2F3 is not a prominent cell type in invasive tissue, consistent with our prior studies in mice[20] and previous reports in human samples[32]. Overall, our findings highlight that tuft-like cell abundance in all IPMN samples trended towards spatial enrichment and that the presence of high-density POU2F3^+^ clusters was obscured by whole tissue quantification, which can only be resolved through spatial analyses.

### Endocrine/Enteroendocrine cells are focally enriched in IPMN

EECs present as solitary cells in pancreatic ducts during neoplastic transformation[17] and development of IPMN and PDAC[18,19,33], which suggests a potential association with disease progression. In addition to the 38 IPMN we performed mxIHC on for broad endocrine marker CHGA, we expanded our cohort by 22 patient samples using IHC. Patient IPMN spanned LG (n=26) and HG (n=27) IPMN to INV (n=7), as well as intestinal (n=8), GF (n=28), and PB (n=24)-type IPMN (**Table S1**). The abundance of CHGA+ IPMN cells did not differ by degree of dysplasia, presence of invasive disease, or histologic subtype **(Figure S1B)**. However, like POU2F3, immunostaining identified CHGA^+^ cell clusters across all subtypes of IPMN (**Figure 3C**). We performed Ripley’s K analysis to determine whether CHGA^+^ IPMN cells exhibited significant clustering and observed that L(obs–theo) showed values above 0 in nearly all samples, indicating clustering across the cohort (**Figure 3D**). Forty-six samples contained enough CHGA^+^ cells for analysis and 35/46 showed significant spatial enrichment by maxL (p <0.05) (**Figure S4A**). Just as for the analysis of POU2F3^+^ IPMN cells, Moran’s I[30] was performed as an orthogonal validation for spatial autocorrelation analysis, and among the analyzed samples, 21/43 exhibited significant positive spatial autocorrelation, with Moran’s I values >0.1 (p<0.05). (**Figure S4B-D**). Moran’s I values ranged from <-0.10 to 0.63. Strong positive spatial autocorrelation (>0.3) was observed in three samples and 17 samples showed significant clustering by both maxL and Moran’s I (**Figure S4D-E**). Collectively, these findings are consistent with the presence of spatially distinct CHGA⁺ endocrine/EEC clusters that are not captured by whole-tissue quantification applications.

### Endocrine clusters include glucagon-expressing alpha cell expansion

We and others have shown that peptide hormone-producing endocrine cells as well as gut EECs populate ADM and pancreatic neoplasia[17,19,34,35]. To determine whether distinct endocrine/EEC subtypes constitute the identified CHGA^+^ clusters, we performed immunofluorescence (IF) staining for insulin (INS, β cells), glucagon (GCG, ɑ cells), pancreatic polypeptide (PP, γ cells), somatostatin (SST, *δ* cells), ghrelin (GHRL, *ε* cells), serotonin (5-HT, enterochromaffin cells), and broad endocrine marker synaptophysin (SYP) in a subset of 20 IPMN (**Figure 4A; S5A**). Among the hormone^+^/SYP^+^ subtypes, GCG and 5-HT were most abundant in both LG and HG IPMN and were concentrated in GF IPMN, comprising up to 80% and 84% of SYP+ IPMN cells, respectively **(Figure 4B-C; S6).** The least abundant hormone^+^ populations across all grades were PP, SST, and GHRL; SST and GHRL were lowly expressed in all subtypes. Of the islet cell types present in IPMN, GCG alone spatially clustered by immunostaining and by Ripley’s K and Moran’s I analyses (**Figure 4D-E**). We observed that 5/6 samples analyzed exhibited significant clustering (p < 0.05) (**Figure 4D; S5B**) while only a single sample showed significantly strong spatial autocorrelation (**Figure 4E**). Consistent with this, NanoString analysis of an independent sample (IPMN Intestinal 2) revealed high expression of GCG in non-islet ROIs (**Figure 1C-D; 4A**), suggesting possible clonal expansion of ɑ-like cells in these tumor lesions. Further, though not typically found in the adult pancreas, some IPMN expressed gastrin (GAST), as previously reported[19,36]. GAST was co-expressed with 5-HT (**Figure S5C**) as in intestinal EECs, and our murine studies in PanIN[17]. Altogether, these analyses show that spatial enrichment of endocrine/EECs in some IPMN lesions includes GCG expressing cells.

**Figure 4.**
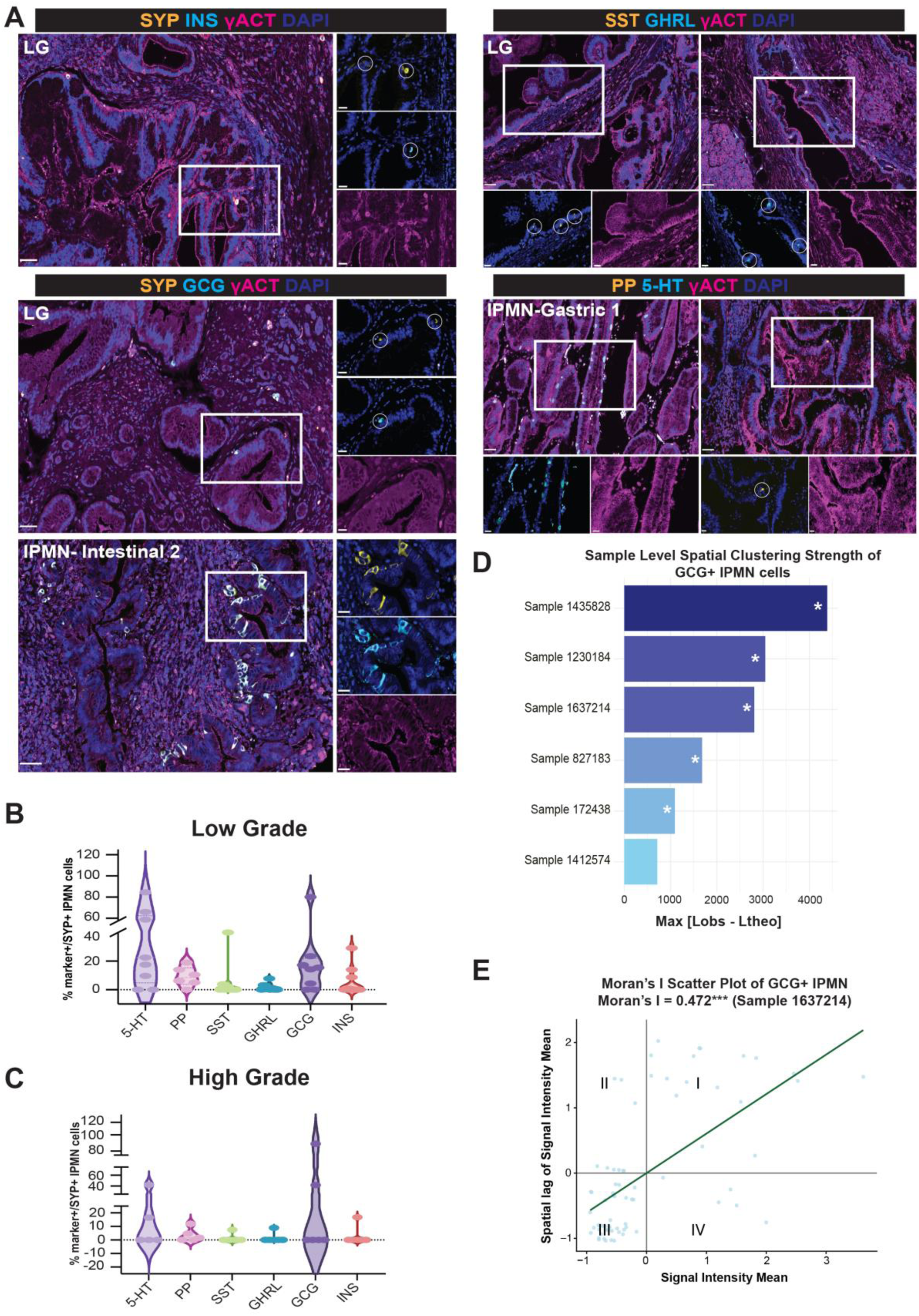
Islet endocrine and enteroendocrine cell subtypes are focally enriched in IPMN lesions. (A) Immunofluorescence (IF) for endocrine markers, synaptophysin (SYP), pancreatic polypeptide (PP), ghrelin (GHRL), glucagon (GCG), somatostatin (SST), insulin (INS), and serotonin (5-HT) in IPMN. Scale bars, 50 μm; insets, 20 μm. White circles, positive stain in single cells. (B) Quantification of each hormone/neurotransmitter in low-grade (LG) and (C) high-grade (HG) IPMN (n=20). (D) Bar plot of clustering strength (Max [Lobs-Ltheo]) of GCG^+^ IPMN cells. Clustering strength was measured using the maximum absolute deviation between observed and theoretical Ripley’s L functions (n=6). (E) Scatter plot of staining intensity of each cell plotted against the average staining intensity of its neighboring cells, termed spatial lag, in one IPMN sample (1637214). The regression slope corresponds to the Global Moran’s I value. Significant positive spatial autocorrelation shown, as a significant positive Moran’s I indicates that cells with similar staining intensity are spatially grouped. Spatial enrichment was defined when Max |Lobs − Ltheo| >95th percentile of a permutation-based null distribution (999 permutations). Wilcoxon rank-sum test with continuity correction. *, p < 0.05 denotes spatial enrichment.

### Enterochromaffin cells are spatially enriched in Low-grade, Gastric IPMN

Enterochromaffin cells were the most abundant EEC population in GF and LG-IPMN epithelium. As 5-HT has been shown to drive tumorigenesis[37], we next quantified 5-HT abundance in a cohort of 21 samples, adding one sample to the initial 20-sample set. For this analysis, we performed IHC as a more sensitive detection method and quantified 5-HT and SYP expression. The abundance of 5-HT^+^ and SYP^+^ IPMN cells were comparable across IPMN grades and subtypes (**Figure 5A-B, S7A-B**) and up to 100% of SYP^+^ cells were 5-HT^+^ in all grades as well as in GF and PB IPMN (**Figure S7C**). Because EECs appeared focally clustered in some samples but more broadly distributed in others (**Figure 5A,C**), we performed Ripley’s L analysis to quantify their spatial organization. Up to 17 samples contained enough EECs for analysis. Ripley’s L showed that most samples had L(obs–theo) values above 0, indicating non-random clustering (**Figure 5D**) and were significantly clustered by MaxL (**Figure S8A**). Further, Global Moran’s I analysis showed significant spatial autocorrelation in 14/16 SYP-stained IPMN and 8/15 5-HT-stained IPMN samples that met the criteria for this analysis. Moran’s I values for SYP ranged from 0.1 (weak) to >0.3 (strong), and three 5-HT samples exhibited values approaching 0.3 (**Figure S8C**). Together, these findings suggest that enterochromaffin cells represent expansion of an EEC-like state within IPMN, adding an additional layer of epithelial heterogeneity beyond overall cell abundance.

**Figure 5.**
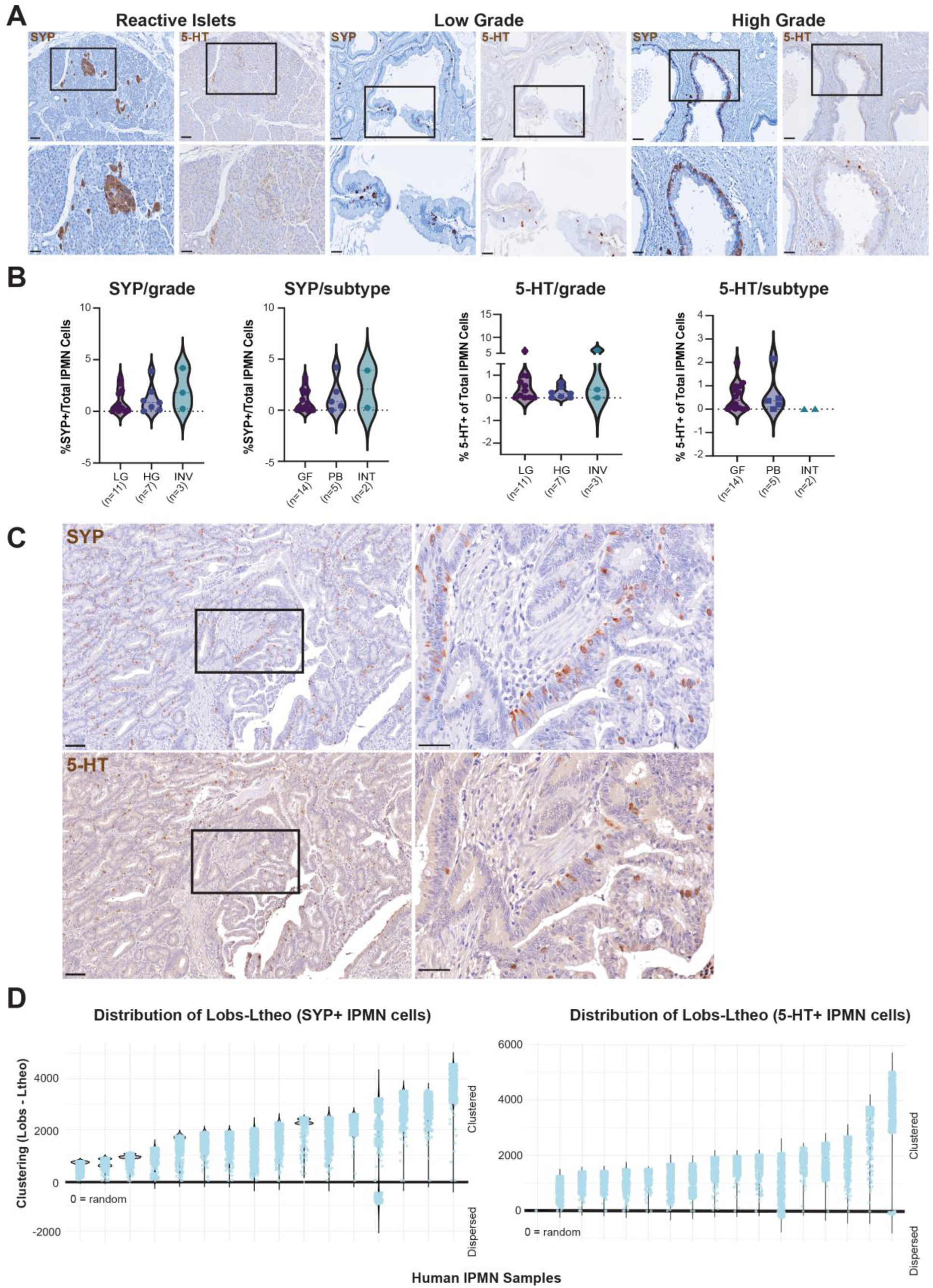
Enterochromaffin cells are focally enriched in IPMN. (A) IHC for SYP and 5-HT in LG and HG IPMN as well as (B) quantification of staining by grade or subtype (n=21). (C) Representative IHC images of SYP^+^ and 5-HT^+^ cells in an INV IPMN sample. (D) Violin plots showing the distribution of Ripley’s L(obs–theo) values across increasing radii for each IPMN sample enriched for SYP (n=16) or 5-HT (n=17). The black line marks 0, where observed clustering equals random expectation; values above or below 0 indicate clustering or dispersion, respectively. Samples are ordered by increasing clustering strength. Scale bars, 100 μm; insets, 50 μm.

### Tuft-like clusters in IPMN are associated with distinct stromal populations

Given that EECs and tuft-like cells show consistent aggregation in human IPMN lesions, we next sought to understand what biological features might be associated with this spatial patterning. To determine what features in the microenvironment of these clusters distinguish them from non-clustered areas, we analyzed spatial transcriptomics data from a four TMA cohort of 31 samples spanning normal, IPMN, and invasive carcinoma using 10x Genomics Xenium *in situ* spatial transcriptomics, generated by the Trinh laboratory[38]. UMAP projection of the integrated dataset across samples shows the major epithelial and stromal (fibroblast/immune/endothelial) populations in this dataset using markers as defined by the original authors **(Figure 6A)**. To determine whether clusters of cells expressing distinct secretory cell markers (e.g., *POU2F3*) were associated spatially with distinct stromal populations, we applied a radius-search algorithm to link stromal cells to adjacent IPMN epithelial cells within a <100μm radius. As in **Figure 3A**, we identified *POU2F3*^+^ cells in all grades of IPMN **(Figure 6B)**. We defined the composition of marker^+^ IPMN tumor cell niches and the surrounding epithelial-adjacent stromal cells in LG, HG and INV tissues. Within the 100μm neighborhood dataset, marker^+^ tumor cells were defined by having ≥ 4 transcripts of *POU2F3,* and clusters were defined by having ≥7 *POU2F3*^+^ tumor cells within a 75μm radius. We found that two samples contained a total of up to 9 *POU2F3^+^* clusters, and that of the fibroblast population, fibroblasts with unresolved subclass identity due to limited Xenium gene panel coverage were significantly enriched near *POU2F3^+^*clusters as compared to negative clusters. Further, and potentially more importantly, activated iCAF (*SOCS3, STAT3*), and myCAF (*FN1, POSTN, TGFB1)* populations were significantly enriched distal to these clusters (**Figure 6C-E**). Of the immune and vascular populations, monocytes (*RGS1*) and resident memory T cells (*CXCR6, ZNF683*) were significantly enriched near *POU2F3*^+^ clusters. Anti-inflammatory macrophages (*CD163*), inflammatory endothelial (*CD74*, *IL1R1*), cytotoxic T (*CD8*) and dendritic (cDC2)(*CD1C, FCER1A*) cells were significantly enriched distal to these positive clusters **(Figure 6F-H)**. These data are consistent with our prior findings in mice showing that POU2F3^+^ tuft cells inhibit stromal activation and suggest that tuft-like niches appear to preferentially associate with stromal and immune cell populations lacking canonical activation markers.

**Figure 6.**
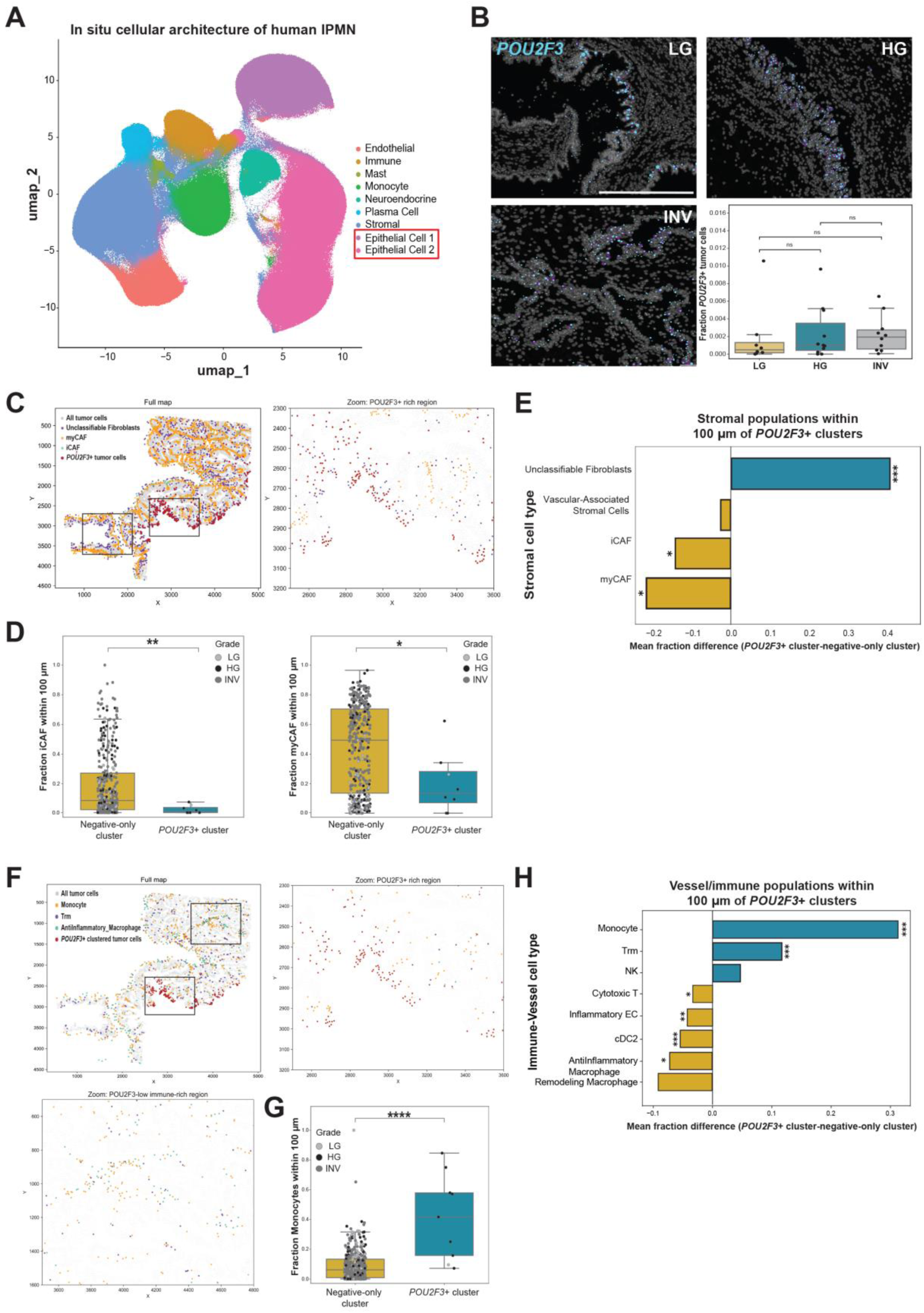
*POU2F3*^+^ IPMN clusters are associated with a defined stromal niche. (A) Uniform Manifold Approximation and Projection (UMAP) showing annotated clusters of integrated epithelial and stromal populations of 31 patient samples spanning normal pancreas, LG, HG IPMN and invasive PDAC from four tissue microarrays (TMAs) generated using 10x Genomics Xenium. (B) Representative images of *POU2F3*^+^ cell clusters in LG, HG and INV samples by transcript, derived from Xenium Explorer 3.2.0. Dot size represents number of transcripts per cell. Box plot shows the fraction of *POU2F3*+ cells among all tumor cells in each sample, stratified by LG (n=8), HG(n=11) and Inv (n=10) IPMN. (C,F) Spatial maps showing selected cell types in a HG IPMN sample and *POU2F3*+ cluster regions. (D,G) Box plots show cluster-level cell type fractions. Each point represents a cluster colored by grade (n=389 negative clusters and n=8 positive clusters in D; n=401 negative clusters; n=9 positive clusters in G). (E, H) Bar plots showing the mean difference in local cell type composition within 100μm of defined *POU2F3*^+^ clusters relative to negative clusters in LG, HG and Inv IPMN (n≤9 clusters). Positive and negative values indicate enrichment near *POU2F3*^+^ and negative clusters, respectively (n=25 samples: 2 with positive clusters and 23 with negative). Trm, tissue resident memory T cell; NK, NK cells; cDC2, conventional dendritic cell type 2. Scale bars, 500 μm for B and coordinates in μm for C,F. All comparisons used two-sided Mann-Whitney U tests; Benjamini–Hochberg FDR correction for bar plots. Asterisks either indicate FDR-adjusted significance(***, FDR < 0.001; **, FDR < 0.01; *, FDR < 0.05) or unadjusted p values ( *,P<0.05, ***,P<0.001, ****,P<0.0001).

## DISCUSSION

PDAC arising from IPMN is characterized by substantial heterogeneity where lesions with similar histological or molecular features may follow different clinical paths[39]. Though IPMN-derived PDAC make up a much smaller percentage of PDAC onset cases (10-15%)[8,40], deciding when to resect these cysts has proven to be difficult, especially in older patients. This suggests that a more accurate assessment of malignant potential will require investigating the underlying biology of disease progression and those factors that drive dysplasia and invasion, providing context to accompany existing criteria.

By integrating NanoString GeoMx and Xenium spatial transcriptomics, immunostaining, quantitative digital pathology analysis (QuPath), and spatial statistics, we show that IPMN are characterized by substantial epithelial heterogeneity, including tuft-like cells and endocrine/EECs. Further, we identified significant spatial clustering of these populations. Whole tissue quantification revealed that the overall abundance of these populations was comparable across all conditions examined in our analysis, masking regional expansion of tuft-like cells and EECs. Spatial statistical analyses, Ripley’s K and Moran’s I, however, demonstrate that positive cells form non-random clusters, suggesting that spatial organization–rather than overall percentage of positive cells—distinguishes tissue architecture across conditions. These findings align with the idea that spatial organization of the tumor microenvironment has a lasting impact on tissue biology and therefore treatment[41]. Our previous work showed that secretory cell types typically absent from the adult pancreas reemerge in response to tissue injury or neoplastic transformation[16,17,21,22]. In the intestines, both tuft cells and EECs arise from a shared SOX4+ secretory progenitor cell[42] and elements of this developmental program reemerge during metaplastic and neoplastic progression in the pancreas[16,34,43].

Tuft cells in the normal human pancreas are found in the intra-and interlobular ducts and are generally absent from the main pancreatic duct and endocrine compartments[44]. Though rare, these cells expand in ADM and neoplasia and are functional, inhibiting pancreatic tumorigenesis through secretion of prostaglandin D_2_ (PGD_2_) in mice[21]. In this study, POU2F3^+^ tuft-like cells were identified in all grades and histological subtypes of IPMN, with some samples exhibiting pronounced spatial clustering. This clustering was not biased towards any grade or subtype, and clusters were surrounded by a potentially tumor-restraining niche as determined by reduced stromal activation. Given that POU2F3^+^ cells emerge in inflammatory contexts and secrete prostaglandins to inhibit malignant progression[45], their presence in HG lesions suggests that multiple mechanisms could be driving their formation and expansion. Perhaps in these HG samples, an amalgamation of inflammatory signals, altered differentiation programs, and broader dysregulation or “confusion” of tumor cell identity is at play[46], resulting in clonal expansion of a protective metaplastic cell state.

EECs are solitary gastrointestinal cells found in normal ducts or in ADM/PanIN and differ from pancreatic islets in hormone composition. Pancreatic ductal and islet cells are developmentally related[47] and share conserved regulatory programs which can be reactivated during neoplastic transformation[34,35], therefore it is unsurprising that we see islet cells in the IPMN ductal epithelium. Moreover, expansion of EECs or endocrine cells (e.g., CHGA, GCG, SST, 5-HT) occurs in other tumors of the pancreas[48], neuroendocrine tumors (PNET), and have been shown in case reports to appear concurrent with IPMN[49]. In our study, we observed that of the islet cell types, only ɑ cells (GCG^+^) exhibited significant focal enrichment across IPMN samples. ɑ cell-rich microadenomas have previously been associated with PanINs[50]. Alterations in amino acid signaling are also associated with ɑ cell expansion[51], where ɑ cells may support regeneration in the diseased pancreas. Therefore, ɑ cells in IPMN may serve to support the resolution of the lesion although this requires further testing. It is tempting to speculate that glucagon receptor targeting therapies for weight loss and diabetes could influence IPMN resolution or progression[52–54] and future cancer rates.

Enterochromaffin cells were the most abundant EEC population with significant spatial clustering in LG and GF IPMN. Enterochromaffin cells have been identified in ADM[17] and are reported to drive disease progression[55]. They are increased in PDAC, where 5-HT signaling through HTR2B supports tumor glycolysis in the context of metabolic stress[37]. Because 5-HT accumulation in the pancreas is often associated with metabolic stress and drives inflammation and fibrosis, the aggregation of 5-HT^+^ cells in IPMN lesions could reflect stressed niches. Alternatively, 5-HT secretion could signal in a paracrine fashion to tissue stroma, driving neoplastic progression.

Endocrine cells within IPMN could arise through several mechanisms. First, neoplastic ductal cells exhibit marked plasticity and, under inflammatory and oncogenic conditions, may aberrantly activate endocrine gene programs[56–58]. Second, progressive neoplastic growth may physically envelop adjacent islets or extend into islets, thereby incorporating endocrine cells into growing lesions. Collectively, these possibilities reflect how disrupted cell identity and spatial organization could impact neoplastic epithelial cell composition. Determining whether the focal enrichment of these cells arises through any these mechanisms will require further investigation.

The cellular heterogeneity and clustering observed in IPMN suggest that these features could be used to refine subtype classifications. At a diagnostic level, however, sampling bias remains challenging due to limited biopsies or fine needle aspirations (FNAs) that could omit endocrine-rich or tuft cell-rich regions, underestimating lesion complexity. Further, there are gaps in how the data from our study can be applied. IPMN samples vary in size and show significant molecular and histological heterogeneity, and we cannot capture the full landscape of an IPMN, where different or mixed-type clonal outgrowth may exist. A larger and more comprehensive spatial dataset could inform diagnosis, risk stratification, and any therapeutic decision making.

This work has several limitations worth noting. For instance, marker expression alone does not confirm that the identified populations are actively secreting effectors or are functional and could, therefore, directly modulate their microenvironment. Further, our NanoString and Xenium analyses were limited to a small number of IPMN samples (e.g., INV, Int), which reduced our ability to detect subtle differences across subtypes and grades. In addition, the Xenium dataset consisted of only 480 genes, which made it difficult to fully subclassify cells, including the fibroblast population we found proximal to *POU2F3*^+^ clusters in IPMN. Further, our whole-tissue quantification did not reveal any significant differences in cell-type abundance for the tuft-like, endocrine, and EEC markers, suggesting that a much larger cohort would be necessary to identify significantly greater overall populations across IPMN subtype and grade. Because IPMN are heterogenous, we annotated each sample based on the highest grade or most abundant subtype. Though practical, this approach may have skewed subtype and grade counts.

In this study, we expand the current understanding of epithelial heterogeneity in IPMN and show that secretory cell populations, such as tuft-like, EECs, and islet endocrine cells, are a common feature in all subtypes and grades. We demonstrate that hormone^+^ cells are selectively expanded in IPMN, with enterochromaffin cells being the most predominate in LG lesions. Both these and glucagon-expressing cells show focal enrichment in IPMN. Finally, we identify an expansion of POU2F3^+^ IPMN cells and identify an associated stromal niche, revealing an additional and previously underappreciated component of heterogeneity within these tumors.

## ACKNOWLEGEMENTS

The authors would like to thank Frank Revetta for sectioning tissue blocks.

## AUTHOR CONTRIBUTIONS STATEMENTS

VQT, KD, and AC conceptualized and planned experiments. MCBT, KD, VQT, and AS secured funding. AC and VQT conducted histopathology while VQT and SS conducted scoring and pathological assessments (grading). AC, BJ and KG performed image quantification analyses and AC conducted spatial statistics. MB and VQT provided spatial neighborhood analysis pipeline and Xenium spatial data. KD supervised the project. EDD provided conceptual feedback. MCBT conducted IPMN resections and provided patient samples. AC and KD wrote the first draft of the manuscript while MCBT, VQT, EDD, KG, BJ, CT and SS reviewed, edited and provided critical feedback on the manuscript. All authors contributed to the final version of the manuscript.

## SUPPLEMENTARY MATERIAL ONLINE

### Supplementary Materials and Methods

**Figure S1.** Secretory cell abundance across IPMN

**Figure S2.** Moran’s I and Ripley’s K analyses demonstrate focal clustering of POU2F3^+^ cells in IPMN

**Figure S3.** POU2F3 IHC scoring and quantification across IPMN grade and subtype

**Figure S4.** Moran’s I and Ripley’s K analyses demonstrate focal clustering of CHGA^+^ cells in IPMN

**Figure S5.** Heterogeneous hormone/neurotransmitter expression patterns in IPMN

**Figure S6.** Hormone/neurotransmitter^+^ cell abundance across IPMN

**Figure S7.** Enterochromaffin cell abundance across IPMN grade and subtype

**Figure S8.** Moran’s I and Ripley’s K analyses demonstrate focal clustering of enterochromaffin cells in IPMN

**Table S1.** Summary of clinicopathologic characteristics of patients from Vanderbilt University Medical Center (VUMC) Cohort

**Table S2.** Primary antibodies used for multiplex immunohistochemistry (mxIHC), immunohistochemistry (IHC), or immunofluorescence (IF) studies

## DATA AVAILABILITY STATEMENT

Clinical data are not publicly available due to inclusion of potentially identifiable and sensitive patient content in accordance with institutional ethical guidelines. De-identified data may be available upon request from the corresponding author and is subject to institutional approval. Spatial transcriptomics data generated from 10X Xenium genomics can be found at Open Science Framework (accession number yd8zh). Any additional data supporting this manuscript will be made available by the corresponding author upon request.

## ETHICS APPROVAL STATEMENT

Patient tissue samples were acquired from the Vanderbilt University Medical Center institutional cohort under institutional review board approval (IRB #110061).

