## Supplementary Figures for "Spatial analysis of Intraductal Papillary Mucinous Neoplasms reveals secretory cell-enriched neighborhoods"

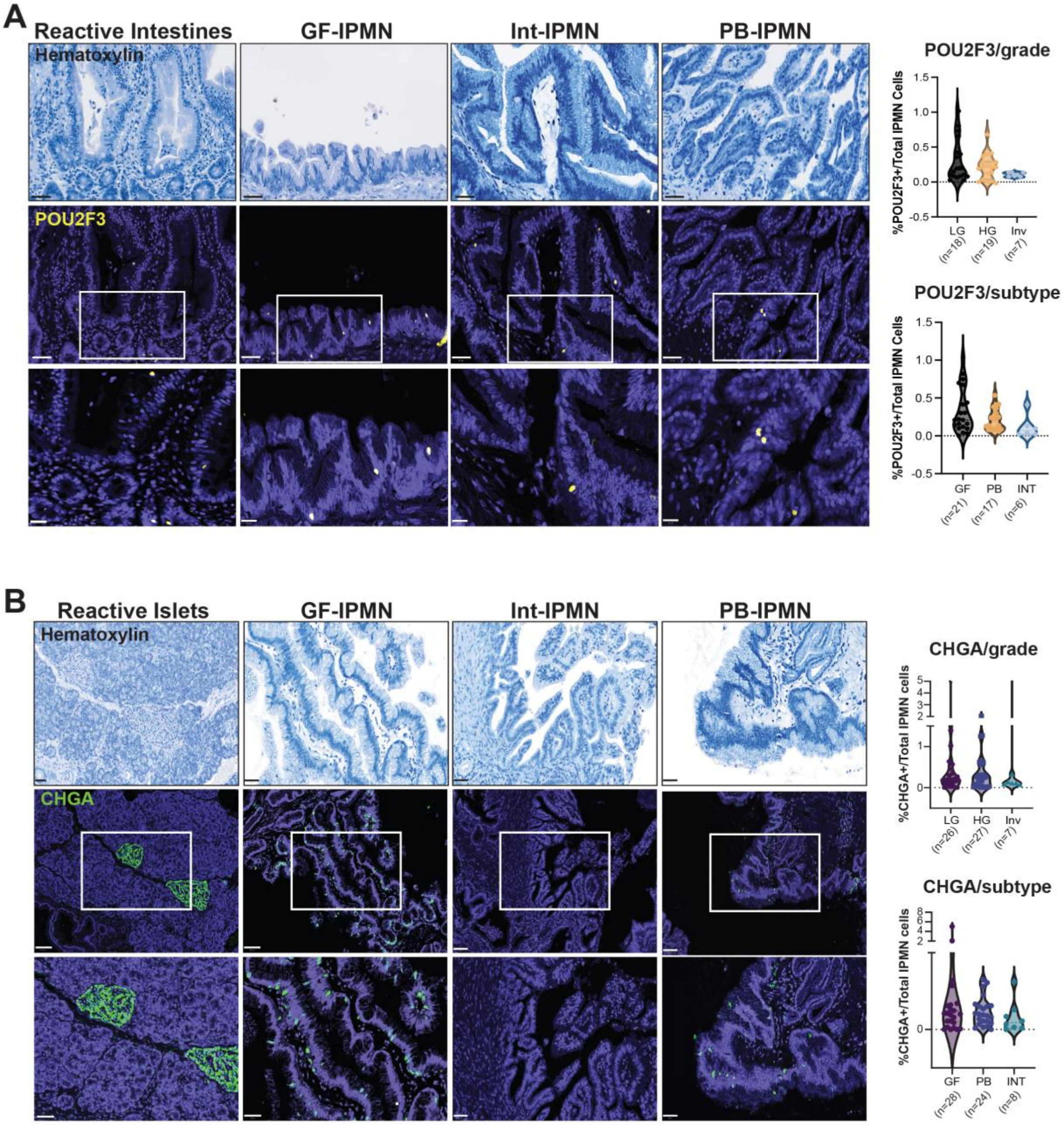


**Figure S1. Secretory cell abundance across IPMN.** (A) Representative images of pseudo-colored POU2F3 IHC (yellow) and hematoxylin (blue) staining in intestines (positive control), GF, Int, and PB IPMN. Quantification of POU2F3$^{+}$ nuclear expression in cells across IPMN samples classified by grade (LG, n=18; HG, n=19, Inv, n=7) and subtype (GF, n=21; PB, n=17; Int, n=6 ). (B) Representative images of CHGA pseudo-colored IHC (green) and hematoxylin staining in islets (positive control), GF, Int, and PB IPMN. Quantification of $CHGA^+$ cell expression across IPMN samples classified by grade (LG, n=26; HG, n=27, Inv, n=7)  or subtype (GF, n=28; PB, n=24; Int, n=8). Top row scale bars, 50 μm; insets, 20 μm.

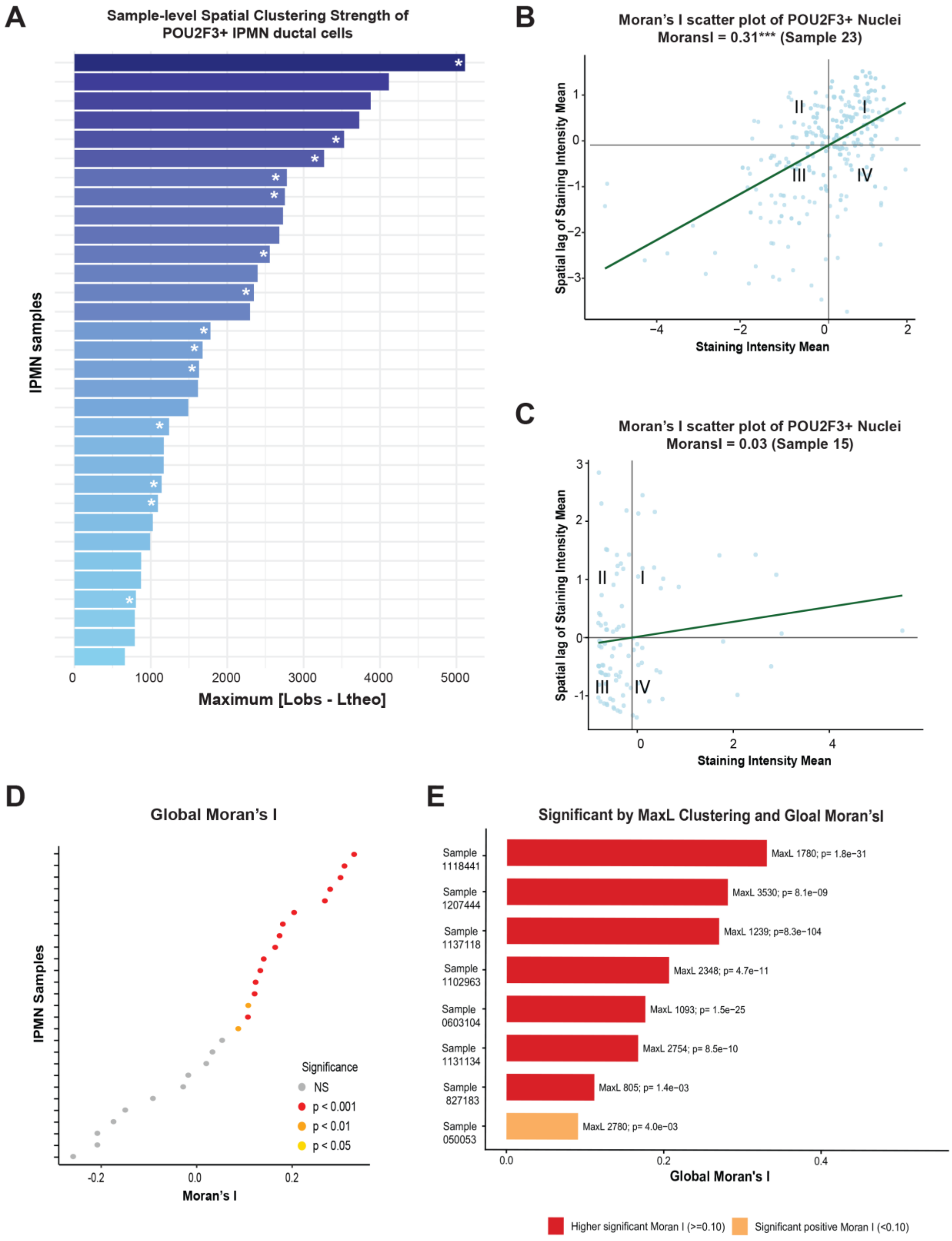
A
Sample-level Spatial Clustering Strength of POU2F3+ IPMN ductal cells
IPMN samples
Maximum [Lobs - Ltheo]
0
1000
2000
3000
4000
5000
B
Moran's I scatter plot of POU2F3+ Nuclei
MoransI = 0.31*** (Sample 23)
Spatial lag of Staining Intensity Mean
Staining Intensity Mean
I
II
III
IV
C
Moran's I scatter plot of POU2F3+ Nuclei
MoransI = 0.03 (Sample 15)
Spatial lag of Staining Intensity Mean
Staining Intensity Mean
D
Global Moran's I
IPMN Samples
Moran's I
Significance
NS
p < 0.001
p < 0.01
p < 0.05
E
Significant by MaxL Clustering and Gloal Moran'sI
Sample 1118441
MaxL 1780; p= 1.8e−31
Sample 1207444
MaxL 3530; p= 8.1e−09
Sample 1137118
MaxL 1239; p=8.3e−104
Sample 1102963
MaxL 2348; p= 4.7e−11
Sample 0603104
MaxL 1093; p= 1.5e−25
Sample 1131134
MaxL 2754; p= 8.5e−10
Sample 827183
MaxL 805; p= 1.4e−03
Sample 050053
MaxL 2780; p= 4.0e−03
Global Moran's I
Higher significant Moran I (>=0.10)
Significant positive Moran I (<0.10)

**Figure S2. Moran's I and Ripley's K analyses demonstrate focal clustering of POU2F3$^+$ cells in IPMN.** (A) Bar plot showing clustering strength (Max [Lobs-Ltheo]) of POU2F3$^+$ nuclei across samples. Each bar represents one sample, and bar length indicates the maximum deviation of Ripley'sL from complete spatial randomness (n=32). Samples are spatially enriched (*, $p < 0.05$) when Max |Lobs − Ltheo| >95th percentile of a permutation-based null distribution (999 permutations). Wilcoxon rank-sum test with continuity correction was performed. (B-C) Moran's I scatter plots: each point represents one cell. The x-axis shows the nuclear staining intensity, and the y-axis shows the average staining intensity of neighboring cells. The regression slope corresponds to Global Moran's I value. Sample 23 (GF, LG) shows significant positive spatial autocorrelation, while Sample 15 (Int, LG) shows weak/non-significant autocorrelation. (D) Global Moran's I plot of p values across IPMN samples. Significance (p values) determined by Z tests (n=27). (E) Samples significant by both MaxL/Ripley clustering and positive Global Moran's I. Bar length shows Moran's I; labels show MaxL deviation and Moran's I p-value (n=8).

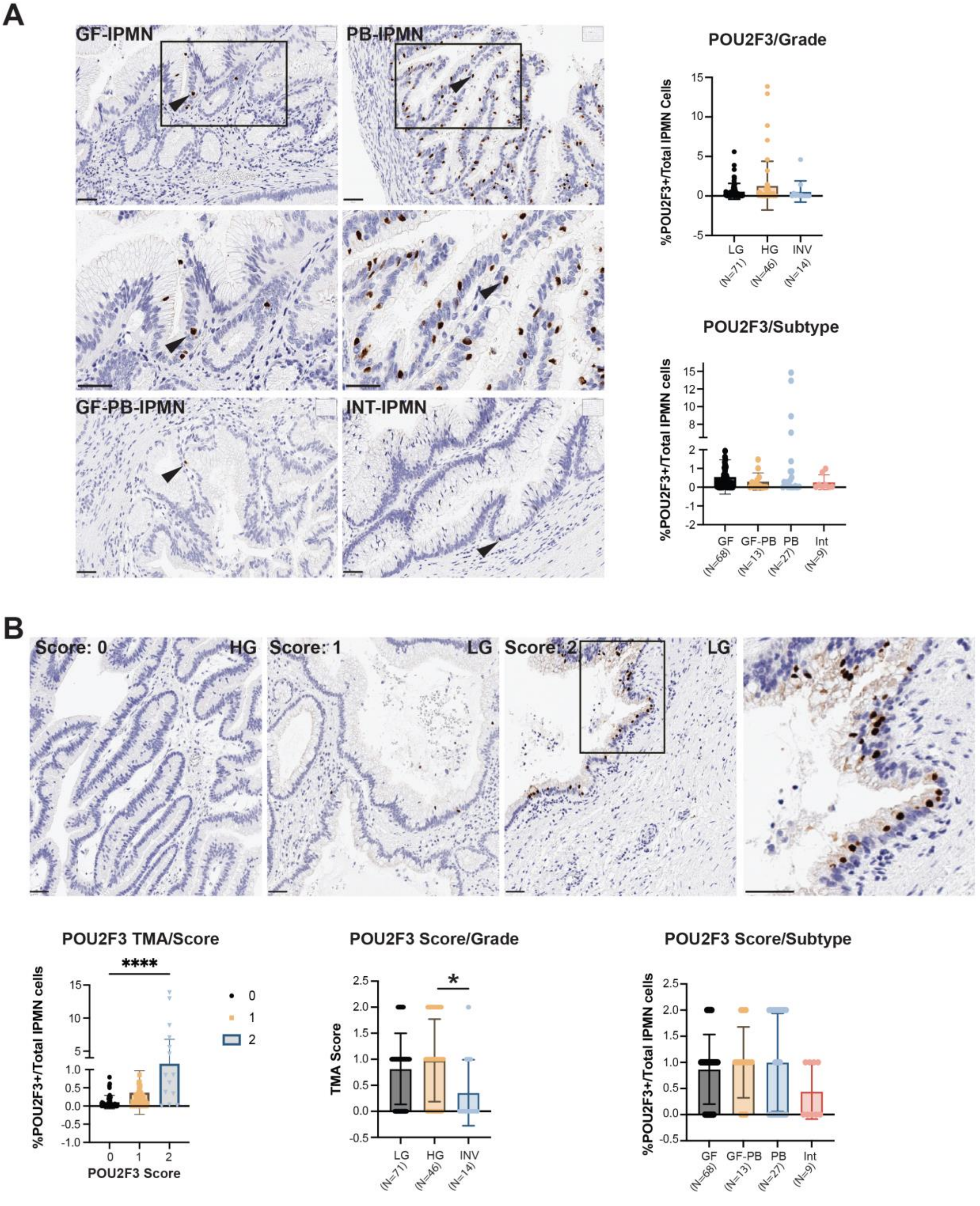


**Figure S3. POU2F3 IHC scoring and quantification across IPMN grade and subtype.** (A) IHC and quantification of POU2F3$^{+}$ cell number as a percentage of total IPMN cells stratified by subtype and grade across IPMN samples. INV samples were excluded from subtype stratification. (B) IHC staining and scoring of POU2F3 expression in tissue microarrays (TMAs) encompassing LG, HG and INV IPMN regions and quantification of % $POU2F3^{+}$ cells by score, stratified by grade (n=131) and subtype (n=117). Scale bars, 50 µm. Brown-Forsythe test followed by One-way or Welch Anova, *, $p < 0.05$; ****, $p < 0.0001$.

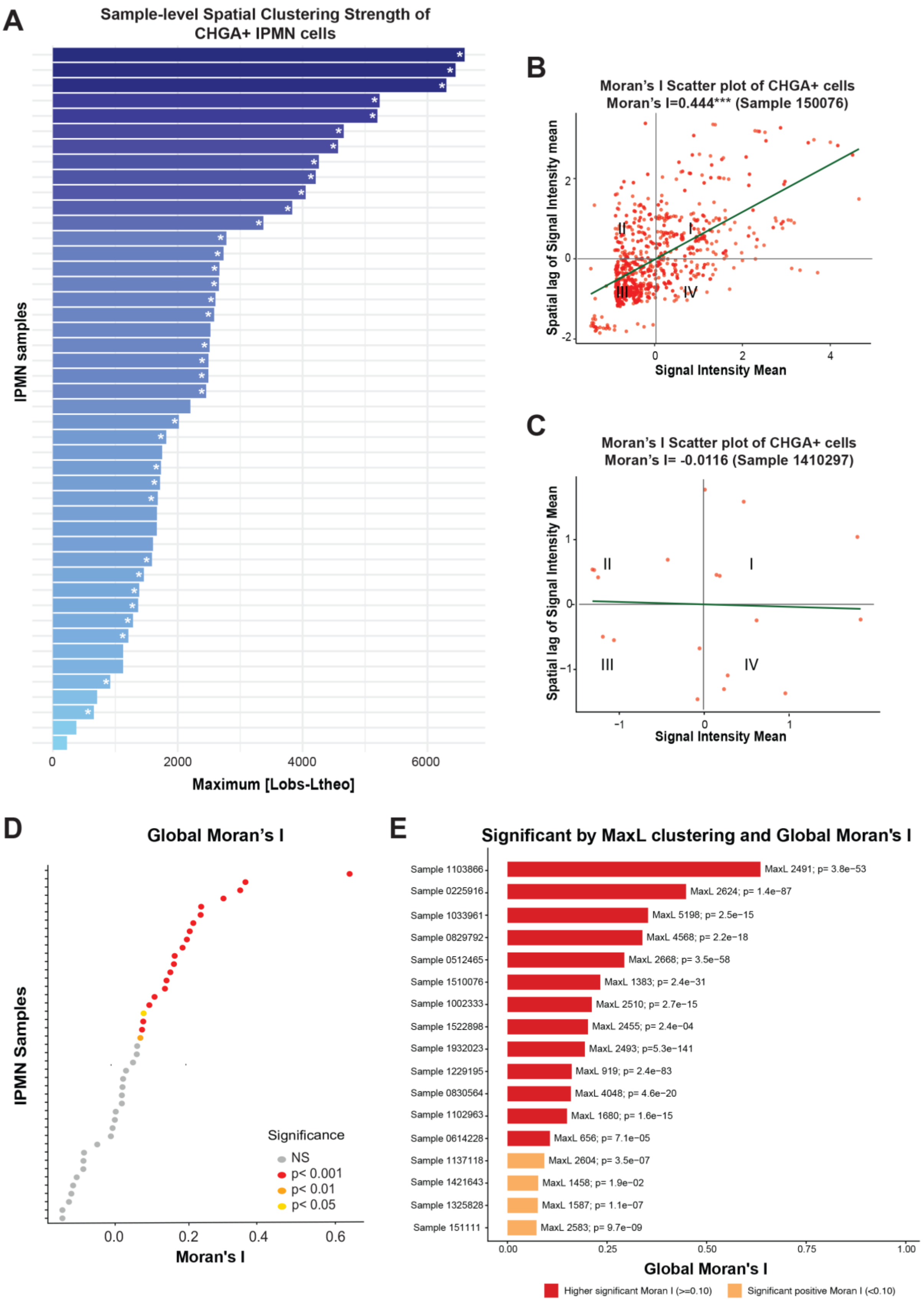
A
Sample-level Spatial Clustering Strength of CHGA+ IPMN cells
IPMN samples
0
2000
4000
6000
Maximum [Lobs-Ltheo]
B
Moran's I Scatter plot of CHGA+ cells
Moran's I=0.444*** (Sample 150076)
Spatial lag of Signal Intensity mean
Signal Intensity Mean
I
II
III
IV
C
Moran's I Scatter plot of CHGA+ cells
Moran's I= -0.0116 (Sample 1410297)
Spatial lag of Signal Intensity Mean
Signal Intensity Mean
I
II
III
IV
D
Global Moran's I
IPMN Samples
Moran's I
Significance
NS
p< 0.001
p< 0.01
p< 0.05
E
Significant by MaxL clustering and Global Moran's I
Sample 1103866 MaxL 2491; p= 3.8e−53
Sample 0225916 MaxL 2624; p= 1.4e−87
Sample 1033961 MaxL 5198; p= 2.5e−15
Sample 0829792 MaxL 4568; p= 2.2e−18
Sample 0512465 MaxL 2668; p= 3.5e−58
Sample 1510076 MaxL 1383; p= 2.4e−31
Sample 1002333 MaxL 2510; p= 2.7e−15
Sample 1522898 MaxL 2455; p= 2.4e−04
Sample 1932023 MaxL 2493; p=5.3e−141
Sample 1229195 MaxL 919; p= 2.4e−83
Sample 0830564 MaxL 4048; p= 4.6e−20
Sample 1102963 MaxL 1680; p= 1.6e−15
Sample 0614228 MaxL 656; p= 7.1e−05
Sample 1137118 MaxL 2604; p= 3.5e−07
Sample 1421643 MaxL 1458; p= 1.9e−02
Sample 1325828 MaxL 1587; p= 1.1e−07
Sample 151111 MaxL 2583; p= 9.7e−09
Global Moran's I
Higher significant Moran I (>=0.10)
Significant positive Moran I (<0.10)

**Figure S4. Moran's I and Ripley's K analyses demonstrate focal clustering of CHGA+ cells in IPMN.** (A) Bar plot showing clustering strength (Max [Lobs-Ltheo]) of $CHGA^+$ cells across samples. Each bar represents one sample, and bar length indicates the maximum deviation of Ripley'sL from complete spatial randomness (n=46). Samples are spatially enriched (*, $p < 0.05$) when Max |Lobs − Ltheo| >95th percentile of a permutation-based null distribution (999 permutations). Wilcoxon rank-sum test with continuity correction was performed. (B-C) Moran's I scatter plots: each point represents one cell. The x-axis reflects staining intensity, and the y-axis is the average intensity of neighboring cells. The regression slope corresponds to Global Moran's I value. Sample 1510076 (HG, GF) shows significant positive spatial autocorrelation, while Sample 1410297 (LG, PB) shows weak/non-significant autocorrelation. (D) Global Moran's I plot of p values across IPMN samples (n=43). Significance (p values) determined by Z tests. (E) Samples reaching significance by both MaxL/Ripley clustering and positive Global Moran's I. Bar length shows Moran's I; labels show MaxL deviation and Moran p-value (n=17).

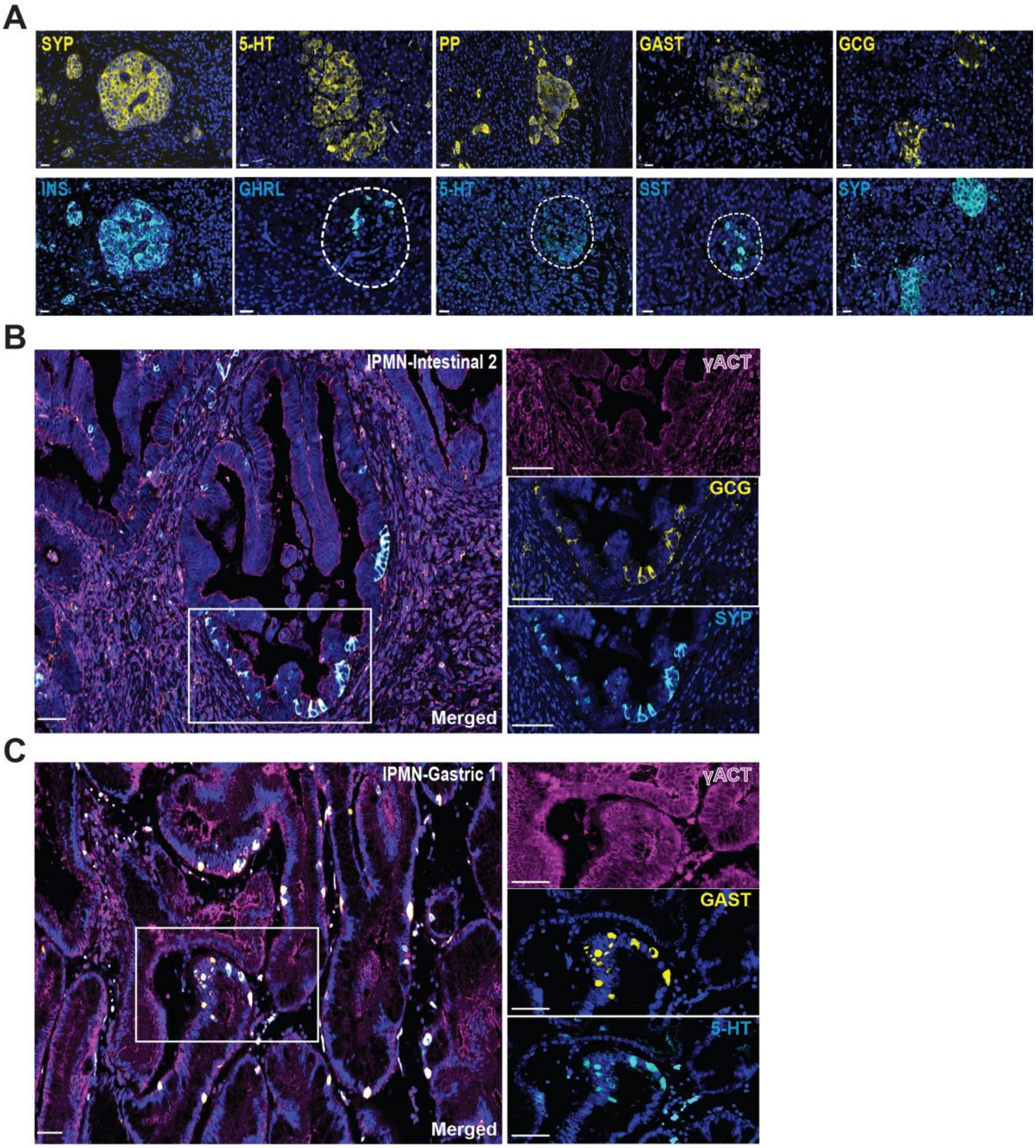


**Figure S5. Heterogeneous hormone/neurotransmitter expression patterns in IPMN.** (A) Immunofluorescence (IF) for broad endocrine marker SYP and endocrine/enteroendocrine subtype cell markers (PP, GHRL, GCG, SST, INS and 5-HT) in pancreatic islets (positive control). (B) Co-

IF for SYP and GCG and (C) 5-HT and GAST within representative IPMN. Scale bars, 20 μm in A and 50 μm in B.

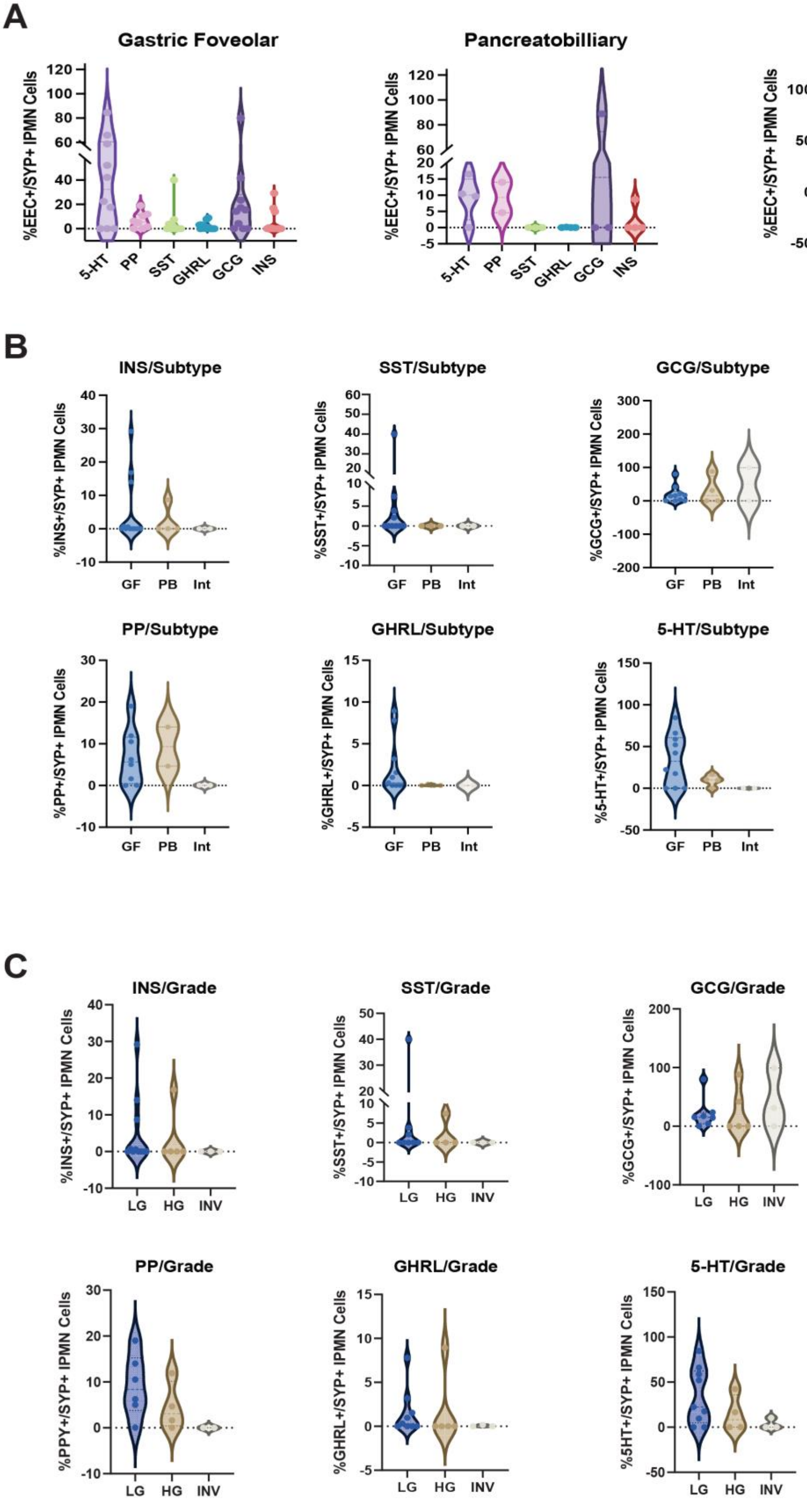
A
Gastric Foveolar
Pancreatobilliary
Intestinal-type
%EEC+/SYP+ IPMN Cells
5-HT
PP
SST
GHRL
GCG
INS
B
INS/Subtype
SST/Subtype
GCG/Subtype
PP/Subtype
GHRL/Subtype
5-HT/Subtype
%INS+/SYP+ IPMN Cells
%SST+/SYP+ IPMN Cells
%GCG+/SYP+ IPMN Cells
%PP+/SYP+ IPMN Cells
%GHRL+/SYP+ IPMN Cells
%5-HT+/SYP+ IPMN Cells
GF
PB
Int
C
INS/Grade
SST/Grade
GCG/Grade
PP/Grade
GHRL/Grade
5-HT/Grade
%INS+/SYP+ IPMN Cells
%SST+/SYP+ IPMN Cells
%GCG+/SYP+ IPMN Cells
%PPY+/SYP+ IPMN Cells
%GHRL+/SYP+ IPMN Cells
%5HT+/SYP+ IPMN Cells
LG
HG
INV

**Figure S6. Hormone/neurotransmitter$^+$ cell abundance across IPMN.** (A) Quantification of endocrine and EEC marker expression (PP, GHRL, GCG, SST, INS and 5-HT) stratified by IPMN subtype. Individual plots of quantification for each marker by IPMN (B) subtype (C) grade (n=20).

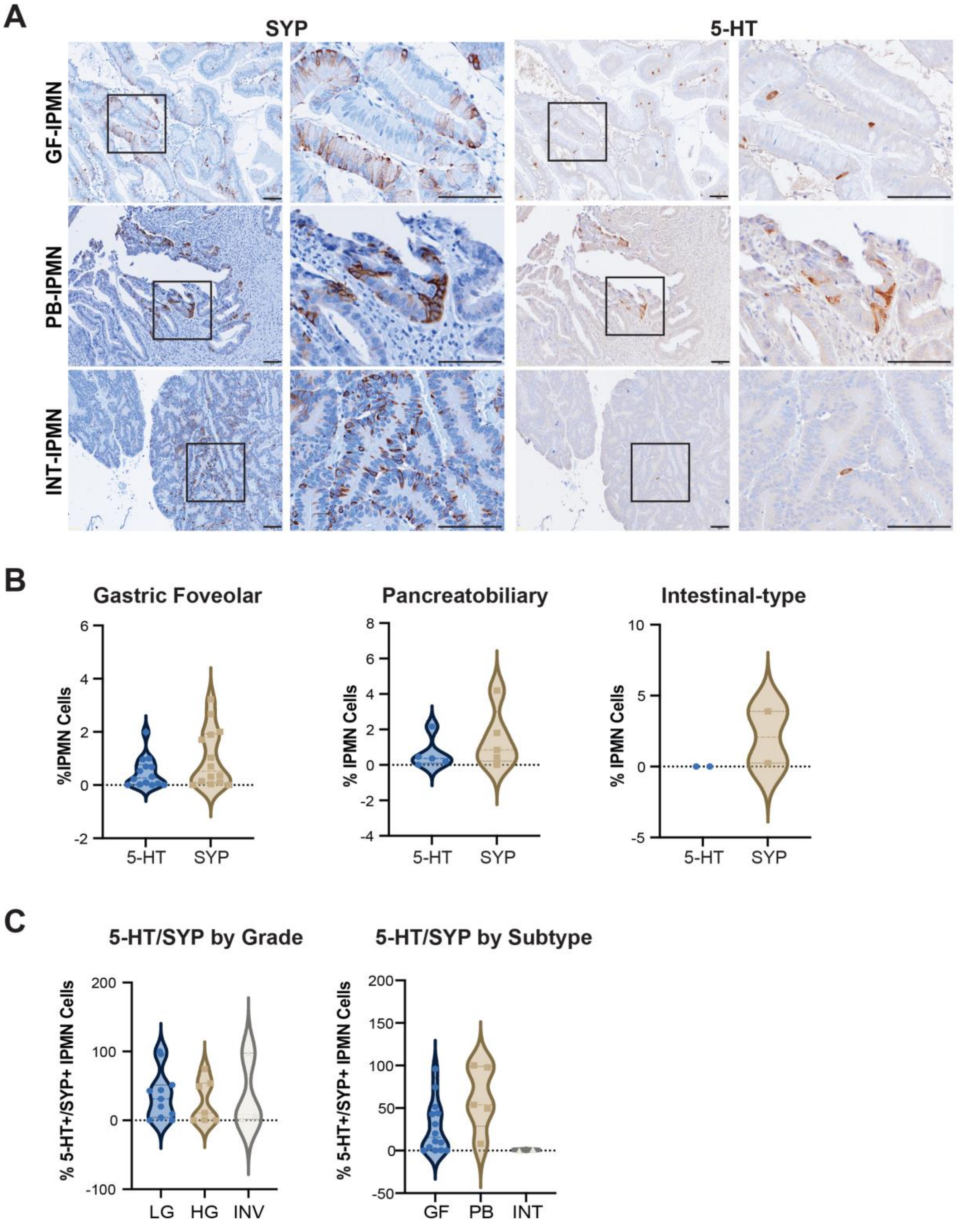
A
SYP
5-HT
GF-IPMN
PB-IPMN
INT-IPMN
B
Gastric Foveolar
Pancreatobiliary
Intestinal-type
%IPMN Cells
% IPMN Cells
5-HT
SYP
C
5-HT/SYP by Grade
5-HT/SYP by Subtype
% 5-HT+/SYP+ IPMN Cells
LG
HG
INV
GF
PB
INT

**Figure S7. Enterochromaffin cell abundance across IPMN grade and subtype.** (A) IHC staining and (B) quantification of SYP and 5-HT in GF, Int, and PB IPMN. Scale bars 100 µm. (C) Percentage of $SYP^+$ IPMN cells that are also $5\text{-}HT^+$ stratified by subtype and grade (n=21).

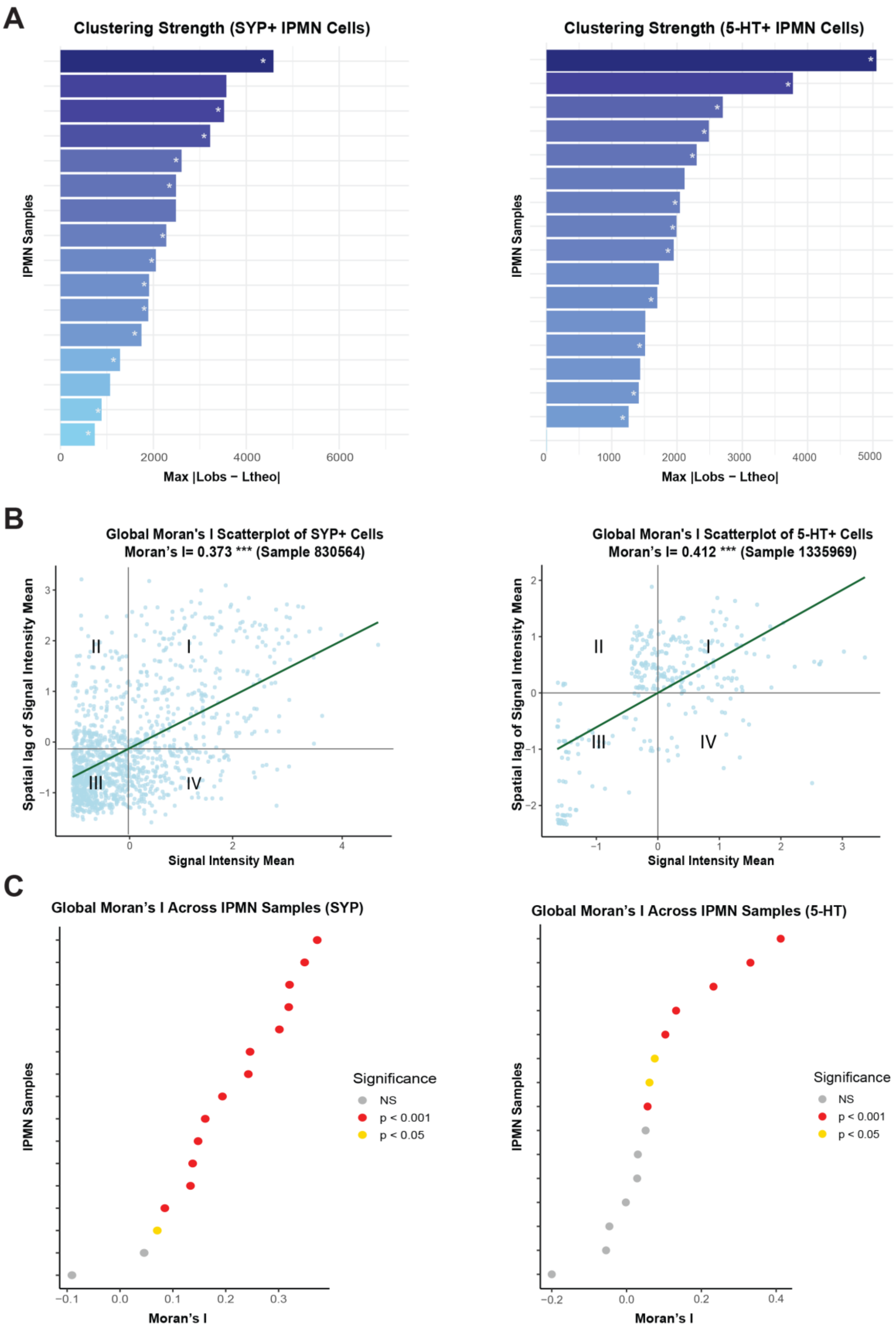
A
Clustering Strength (SYP+ IPMN Cells)
IPMN Samples
Max |Lobs − Ltheo|
0
2000
4000
6000
Clustering Strength (5-HT+ IPMN Cells)
IPMN Samples
Max |Lobs − Ltheo|
0
1000
2000
3000
4000
5000
B
Global Moran's I Scatterplot of SYP+ Cells
Moran's I= 0.373 *** (Sample 830564)
Spatial lag of Signal Intensity Mean
Signal Intensity Mean
I
II
III
IV
Global Moran's I Scatterplot of 5-HT+ Cells
Moran's I= 0.412 *** (Sample 1335969)
Spatial lag of Signal Intensity Mean
Signal Intensity Mean
I
II
III
IV
C
Global Moran's I Across IPMN Samples (SYP)
IPMN Samples
Moran's I
−0.1
0.0
0.1
0.2
0.3
Significance
NS
p < 0.001
p < 0.05
Global Moran's I Across IPMN Samples (5-HT)
IPMN Samples
Moran's I
−0.2
0.0
0.2
0.4
Significance
NS
p < 0.001
p < 0.05

**Figure S8. Moran's I and Ripley's K analyses demonstrate focal clustering of enterochromaffin cells in IPMN.** (A) Bar plots showing clustering strength (Max [Lobs-Ltheo]) of $SYP^+$ and $5\text{-}HT^+$ cells across samples. Clustering strength was measured using the maximum absolute deviation between observed and theoretical Ripley's L functions. Samples were classified as spatially enriched when Max |Lobs − Ltheo| >95th percentile of a permutation-based null distribution (999 permutations). Wilcoxon rank-sum test with continuity correction. *, $p < 0.05$ denotes spatial enrichment (n=16-17). (B) Moran's I scatter plots: each point represents one cell. The x-axis reflects staining intensity, and the y-axis shows the average staining intensity of neighboring cells. The regression slope corresponds to Global Moran's I value. Samples 830564 (LG, GF) and 1335969 (LG, GF) show significant positive spatial autocorrelation for SYP and 5-HT, respectively. (C) Plots of Global Moran's I p values across IPMN samples (n=15-16). Significance (p values) determined by Z tests.
