## Supplementary Tables for "Spatial analysis of Intraductal Papillary Mucinous Neoplasms reveals secretory cell-enriched neighborhoods"

| | Study Population N (%) | Low−grade IPMN N (%) | High−grade IPMN N (%) |
|---|---|---|---|
| **Sample Size** | **60** | **38** | **22** |
| **Age Group (years)** | | | |
| <50 | 1 (1.7) | 0 (0) | 1 (4.5) |
| 50−59 | 13 (21.7) | 8 (21.1) | 5 (22.7) |
| 60−69 | 21 (35) | 11 (28.9) | 10 (45.5) |
| 70−79 | 20 (33.3) | 16 (42.1) | 4 (18.2) |
| 80+ | 5 (8.3) | 3 (7.9) | 2 (9.1) |
| **Sex** | | | |
| Male | 28 (46.7) | 17 (44.7) | 11 (50) |
| Female | 32 (53.3) | 21 (55.3) | 11 (50) |
| **Race** | | | |
| White | 55 (91.7) | 35 (92.1) | 20 (90.9) |
| Black | 4 (6.7) | 2 (5.3) | 2 (9.1) |
| White, Native Hawaiian or Other Pacific Islander | 1 (1.7) | 1 (2.6) | 0 (0) |
| **Whole−IPMN Variant/Subtype** | | | |
| GF | 23 (38.3) | 20 (52.6) | 3 (13.6) |
| PB | 13 (21.7) | 8 (21.1) | 5 (22.7) |
| INT | 9 (15) | 6 (15.8) | 3 (13.6) |
| Mixed subtype | 2 (3.3) | 1 (2.6) | 1 (4.5) |
| IPMN, unspecified subtype | 7 (11.7) | 2 (5.3) | 5 (22.7) |
| Malignant/carcinoma descriptor | 6 (10) | 1 (2.6) | 5 (22.7) |
| **Malignancy Category** | | | |
| Non−malignant | 14 (23.3) | 7 (18.4) | 7 (31.8) |
| Borderline malignant potential | 2 (3.3) | 2 (5.3) | 0 (0) |
| Malignant/invasive carcinoma | 14 (23.3) | 4 (10.5) | 10 (45.5) |
| IPMN, malignancy not specified | 25 (41.7) | 20 (52.6) | 5 (22.7) |
| Retention cyst/uncertain | 5 (8.3) | 5 (13.2) | 0 (0) |
| **Surgical Procedure** | | | |
| Whipple | 40 (66.7) | 27 (71.1) | 13 (59.1) |
| Pancreatectomy | 19 (31.7) | 11 (28.9) | 8 (36.4) |
| Puestow | 1 (1.7) | 0 (0) | 1 (4.5) |
| **History of Diabetes** | | | |
| No | 38 (63.3) | 26 (68.4) | 12 (54.5) |
| Yes | 22 (36.7) | 12 (31.6) | 10 (45.5) |
| **IPMN Location** | | | |
| Main duct | 6 (10) | 3 (7.9) | 3 (13.6) |
| Branch/side branch | 5 (8.3) | 4 (10.5) | 1 (4.5) |
| Unknown/not recorded | 49 (81.7) | 31 (81.6) | 18 (81.8) |

Counts are shown as N (%) within each column. Variant/subtype reflects the whole−IPMN annotation in Book3.xlsx.

**Table S1.** Summary of clinicopathologic characteristics of patients from Vanderbilt University Medical Center (VUMC) Cohort

### Antibodies Used for Immunohistochemistry and Immunofluorescence

| Antibody | Company | Catalog # | Dilution, IHC | Dilution, IF |
|---|---|---|---|---|
| CHGA (rabbit) | Abcam | ab254557 | 1:1,000 | 1:500 |
| Gastrin (rabbit) | BioGenix | PU019−UP | 1:100 | 1:100 |
| Ghrelin (rabbit) | Cell Signaling | 31865 | N/A | 1:1,000 |
| Glucagon (mouse) | Sigma | G2654 | N/A | 1:500 |
| g Actin (488) | Santa Cruz | Sc−65638 | N/A | 1:100 |
| g Actin (546) | Santa Cruz | Sc−65638 | N/A | 1:100 |
| Insulin (Guinea pig) | Fitzgerald | 20−IP35 | N/A | 1:1,000 |
| Pancreatic Polypeptide (rabbit) | Abcam | ab255827 | N/A | 1:80,000 |
| POU2F3 (rabbit) | Sigma | HPA019652 | 1:200 | N/A |
| POU2F3 (rabbit) | Cell Signaling | #Q9UKI9 | 1:500 | N/A |
| Serotonin (goat) | Immunostar | 20079 | 1:20,000 | 1:20,000 |
| Serotonin (rat) | Santa Cruz | Sc−58031 | 1:500 | 1:500 |
| Somatostatin (rat) | Millipore | MAB354 | N/A | 1:1,000 |
| Somatostatin (rabbit) | Abcam | ab111912 | N/A | 1:2,000 |
| Synaptophysin (rabbit) | Cell Marque | 336R−94 | 1:500 | 1:500 |

Dilutions are shown for immunohistochemistry (IHC) and immunofluorescence (IF); N/A indicates not used for that assay.

**Table S2.** Primary antibodies used for multiplex immunohistochemistry (mxIHC), immunohistochemistry (IHC), or immunofluorescence (IF) studies.
