## Supplementary Methods for "Spatial analysis of Intraductal Papillary Mucinous Neoplasms reveals secretory cell-enriched neighborhoods"

### *Multiplex Immunohistochemistry*

Formalin-fixed paraffin-embedded tissues from the VUMC cohort were sectioned at 4 µm and stained as previously described[1]. IPMN samples from VUMC cohort were immunostained for tuft, EECs, and the endocrine cells. Slides were heated to 60°C for 30 min, deparaffinized in xylenes, and rehydrated in an ethanol gradient. Slides were stained with Mayer's hematoxylin, coverslipped with 30% glycerol in PBS and scanned with an Olympus VS200 slide scanner. Subsequently, they underwent antigen retrieval in 10 mM sodium citrate pH 6.0 with a microwave set at maximum power until boiling, reduced to minimum power for 20 min, and left at RT for 30 min. Slides were treated with 3% hydrogen peroxide for 10 min, blocked with Protein Block Serum-Free (Dako) for 10 min, and incubated with the primary antibody overnight. Secondary antibodies were incubated for 1 hour, and the signal was revealed with AEC$^+$ High Sensitivity Substrate Chromogen. The slides were then coverslipped with 30% glycerol in PBS, and were scanned with the Olympus VS200. Subsequently, they were de-coverslipped in PBS (by immersion) and underwent double-distilled water, 70% ethanol, 95% ethanol, ethanol, double-distilled water sequence for 2 minutes each to eliminate the 3-amino-9-ethylcarbazole (AEC) chromogen. The previous sequence was then restarted at the antigen retrieval step until all primary antibodies (**Table S2**) were added on all slides. Scans were loaded in QuPath v0.5.1[2] and registered with the image-combiner v0.3.0 package.

### *Standard Histological Staining*

Sections were deparaffinized in xylenes, rehydrated in a series of graded ethanols, and then washed in PBS-T and PBS. Endogenous peroxidase activity was blocked with a 1:50 solution of 30%

$H_2O_2$:PBS followed by antigen retrieval either by microwave in 100 mM sodium citrate, pH 6.0 or with pH 6.0 citrate buffer in a pressure cooker at 105°C for 15 min, with a 10-min cool down. Sections were blocked with 1% bovine serum albumin and 5% normal goat/rabbit serum in 10mMTris (pH 7.4), 100 mM $MgCl_2$, and 0.5% Tween 20 for 1hr at room temperature. Primary antibodies (**Table S2**) were diluted in blocking solution at room temperature overnight. Slides were washed, incubated in streptavidin-conjugated secondaries (ImmPRESS IgG polymer Kit, Peroxidase; Vector Laboratories, Inc. Burlingame, CA, US) or Dako EnVision+ System-HRP labeled Polymer, and developed with DAB (ImmPACT DAB Substrate Kit, Peroxidase; Vector Laboratories, Inc. Burlingame, CA, US). Tissues were counterstained with Hematoxylin Gill III (3801540; Leica) and cover slipped with resin mounting media, Cytoseal 60 (8310-16; Epredia). For IF, sections were blocked with 10% donkey serum in PBS for 1hr at room temperature and primary antibodies were diluted in primary antibody solution (1% BSA in PBS/0.1% Triton X-100) overnight. Secondary antibodies (**Table S2**) from (Thermo Fischer Scientific, Waltham, MA, USA) were used at 1:500 and incubated at room temperature in a humidity chamber for 1 hour. Tissues were cover slipped using antifade reagent with DAPI (P36931; Invitrogen). Slides were scanned at 20x magnification with an Olympus VS200 slide scanner (Tokyo, Japan). For TMAs derived from the University of Pittsburgh, slides were stained using a Ventana BenchMark Ultra autostainer.

***Cell Quantification and Scoring***

For immunostaining quantification, each tissue section/slide was reviewed by a board-certified GI pathologist (VQT), and grade and subtypes were assigned accordingly. VUMC patient demographic variables in addition to whole-lesion level grading and subtyping of IPMN are reported separately in the main demographic table (**Table S1**). Because IPMNs can be very heterogeneous, the subtype or grade represented on the section/slide may differ from the overall subtype assigned to the patient's IPMN in the demographic table. Samples used for immunostaining included 60 unique analyzed samples spanning the spectrum of disease progression, including low-grade dysplasia (LG; n = 24), high-grade dysplasia (HG; n = 28), and invasive carcinoma (Inv; n=8). Histologic subtypes included intestinal (n = 6), gastric foveolar (GF; n = 28), pancreatobiliary (PB; n = 24) and mixed (n=2) IPMN.

Cells in tissue section slides derived from TMAs and VUMC cohort samples were counted using QuPath. Analysis of mxIHC and IHC samples was carried out on AEC-stained serial sections for POU2F3 and Chromogranin A (CHGA) and on DAB-stained sections for POU2F3, CHGA, synaptophysin (SYP) and serotonin (5-HT). For mxIHC, hematoxylin-stained images were overlaid on either POU2F3 or CHGA AEC-stained images. Cells within IPMN lesions were detected using the Cell Detection function. IPMN ductal epithelium was manually annotated based on cell morphology and classified as "duct" using the Classifier function. All stromal and/or normal acinar tissue areas were excluded. AEC-positive ductal cells were identified using a trained classifier. For IHC samples, the Positive Cell Detection function in QuPath was used (**Fig. 2A**).

POU2F3$^{+}$ cells from TMAs were detected using the Positive Cell Detection function in QuPath. IPMN ductal epithelium—regardless of grade—was classified as "duct", and a threshold classifier for POU2F3-positive cells was applied. POU2F3 immunostaining for low-grade, high-grade, and invasive lesions was scored using a three-point semi-quantitative system (scores 0-2). Only the epithelial elements of the lesions were scored. Score 0 staining pattern corresponded to absent nuclear POU2F3 staining, score 1 corresponded to isolated cells with nuclear POU2F3 staining, and score 2 corresponded to small groups and clusters of cells with nuclear POU2F3 staining.

Immunofluorescence (IF) was performed for γ-actin, DAPI, and the following hormones/neurotransmitters (NT) of interest per slide including, glucagon (GCG, n=16), somatostatin (SST, n=15), (SYP, n=16), gastrin (GAST, n=19), ghrelin (GHRL, n=16), pancreatic polypeptide (PP, n=11), insulin (INS, n=16), serotonin (5-HT, n=17). Annotations, cell detection, and appropriate cell classification were performed as previously mentioned. Some samples did not express all hormones and were not analyzed. Using serial sections, we measured the proportion of cells positive for each hormone/NT relative to total $SYP^+$ IPMN cells. GAST was quantified relative to total $IPMN^+$ cells, as this panel was applied to non-serial tissue sections.

***NanoString Spatial Transcriptomic Profiling***

The NanoString GeoMx Whole Transcriptome Atlas (WTA) platform was used to profile 95 selected regions of interest (ROI) from four IPMN tissue samples from the VUMC cohort. Pathologist-reviewed regions of interest were selected from each IPMN sample for NanoString spatial transcriptomic profiling. Samples S19-32023 (IPMN-Intestinal 1) and S14-21643 (IPMN-Intestinal 2) were both graded as HG with invasive areas and as INT, while S15-10076 (IPMN-Gastric 2) and S11-37118 (IPMN-Gastric 1) were graded as LG and GF. 23-24 ROIs were selected per sample and annotated based on tissue type as either islets/acinar tissue (n=13), IPMN (n=80) ducts glands (n=1) or stroma (n=1). The median nuclei count per ROI is 1,460 and the Interquartile range (IQR) was 1,159 to 2,001. Next Generation Sequencing generated a median of 2,733,631 aligned deduplicated reads per ROI (IQR: 2,047,304 to 5,240,097). Raw count matrices and ROI properties were imported into R from the NanoString GeoMx DSP analysis software. Data analyses were executed using R (version 4.4.2, 2024-10-31 ucrt), tailored for a 64-bit windows platform. R integrated package utilized in the study consisted of dplyr('1.1.4')[3], DESeq2('1.46.0')[4], pheatmap('1.0.13')[5] and matrixStats('1.5.0')[6]. Gene names and ROI identifiers were standardized to consistency for downstream analysis. Count data was normalized using DESeq2. Variance stabilizing transformation was performed to visualize data structure. PCA was performed on VST-normalized data to visualize ROI clustering based on IPMN sub-type. Hierarchical gene cluster was performed using pheatmap. Gene expression values were scaled by z-score and the top 50 genes with the highest variances across all ROIs were selected for visualization.

***Spatial Statistics***

Spatial statistics were performed on IPMN samples from the VUMC cohort immunostained for tuft, EECs, and alpha cells. Whole-slide image files (VSI) were analyzed using QuPath (v0.5.1)[2]. Cellular spatial autocorrelation and clustering were analyzed using Moran'sI[7] and Ripley'sK[8-10], respectively in R. QuPath was used to obtain single-cell centroid coordinates (X, Y; µm), cell classifications, and intensity measurements derived from DAB/AEC or Cy5 staining. All downstream spatial analyses were performed in R (v4.3.1).

Global spatial clustering of cell staining intensity was assessed using Moran's I statistics[11] to determine whether cell intensity values at neighboring locations were correlated (autocorrelation)[12]. Only marker$^{+}$ IPMN ductal cells with quantified DAB/AEC intensity measurements were included in the analysis. Intensity was measured as either nuclear mean

(NucDABMean) for nuclear stains or cellular mean (CellDABMean) for cytoplasmic stains, and global Moran's I was calculated for both values to determine the global spatial autocorrelation within each tissue sample. Samples containing <5 qualifying cells were omitted. Analyses were performed using the spdep R package[13-15]. Spatial relationships between cells were defined using a k-nearest neighbor (kNN) approach based on distance between cell centers. For each sample, the number of neighbors was set to $k = 8$; $k = n - 1$ was used for samples with fewer than nine cells. A spatial-weights matrix was used to define the neighboring cells for every cell. For each cell, the eight nearest cells were defined as neighbors and given equal weights. The weights were scaled to sum to one for each cell (style = "W"), allowing comparisons across tissue samples of different sizes. [11,13,15]. For each sample, Global Moran's I was calculated based on cell or nuclear DAB intensity using the weights matrix based on the eight nearest neighboring cells. The observed Moran's I values, expected values (under spatial randomness), variance, Z-scores, and p-values were obtained using the Moran's I test. Statistical significance was defined as $p < 0.05$. Positive Moran's I values indicate clustering of cells with similar intensities, whereas negative values indicate dispersion. Values near zero indicate random spatial patterning. Results were summarized for all samples and visualized using ranked Morans plots, whereby standardized staining intensities were plotted against the average staining intensities of neighboring cells (spatially lagged values).

To determine whether marker$^+$ IPMN ductal cells were spatially clustered beyond what would be expected by chance, Ripley's K spatial analyses[8-10] was performed. Ripley's K analysis is a distance-based method that determines how well points (cells) cluster across a sample by measuring how many nearby cells are found as the distance around each cell increases (spatial scales). For each sample, Ripley's K(r) calculations were performed within a spatial window encompassing all detected cells, with edge correction used to account for cells near edges. Ripley's K(r) was converted into the L-function, [$L(r)=\sqrt{K(r)/\pi}-r$], which allows for direct comparison of the observed spatial patterning to the expected pattern under random spatial distribution ($L(r) = 0$). [16].To quantify clustering, the difference between the observed and expected L-functions ($L_{obs}-L_{theo}$) across distances was calculated. Positive L(r) values indicate clustering and negative values indicated dispersion. The maximum deviation of this difference, maxL, was used as a single measurement of clustering strength for each sample. To determine if the observed clustering was beyond random, we performed a permutation-based null model[17,18]. In other words, for each sample, the same number of marker$^{+}$ IPMN ductal cells were randomly chosen from all detected cell locations within each tissue, and Ripley's K was recalculated 999 times (999 permutations). This allows us to retain tissue structure and overall cell density while randomizing cell identity. Samples were considered spatially enriched if their observed clustering was above the 95th percentile of the permutation distribution, and p-values were calculated. Statistical significance was defined as $p < 0.05$. Portions of the data processing and visualization code were developed with help from an AI-based language model (ChatGPT, OpenAI), with all code reviewed, edited, and validated by the authors.

### *10x Genomics Xenium in Situ Spatial Analysis*

We reprocessed 10x Genomics Xenium data from four TMAs consisting of 31 human pancreas samples from the Open Research Funders Group (ORF) repository [19]. Cases spanned the spectrum of disease progression, including normal, ADM, LG, HG and INV. We processed data starting from their single-cell spatially resolved and cell-type annotated dataset. A radius-search algorithm[19] was utilized to link stromal cells to epithelial cells and identify the epithelial niche composition of each epithelial-adjacent stromal cell in LG, HG and INV tissues. The algorithm relies on the brute-force calculation of the Euclidean distance between each cell. Stromal, immune, and endothelial cells were queried one by one, calculating their distance to every epithelial cell, to identify those located within 100 μm of queried cell. Output files from the radius-search algorithm contain all stromal-epithelial cell pairs identified within 100 μm distance. Downstream analysis was run using a custom Python 3 script to determine, for each fibroblast and immune/endothelial cell, the number of neighboring IPMN cells expressing genes of interest. Because each tumor cell could appear in multiple rows based on its spatial relationship, a temporary file containing one row per unique tumor cell was created for spatial clustering. The original spatial relationship tables were retained unchanged.

$POU2F3^+$ IPMN tumor cells were identified using a transcript-count threshold of ≥4. Positive tumor cell clusters were identified within each sample using density-based spatial clustering (DBSCAN)[20]. Clusters required ≥7 unique tumor cells within a 75 μm neighborhood radius. Cells not assigned to a cluster were excluded from cluster-level analyses. Clusters identified were merged back into the original spatial relationship tables, and stromal and immune analyses were performed. For each cluster, stromal or immune cells were identified, and the cell-type fractions were calculated as the total number of cells of each type overall all target cells surrounding a cluster. Fractions of each stromal/immune/endothelial subpopulations were compared between $marker^+$ and negative clusters. Analyses were performed in Python using pandas[21], NumPy[22], SciPy[23], matplotlib[24], and seaborn[25]. CSV processing and summary table generation were performed with pandas; numerical operations with NumPy; statistical testing with SciPy; and visualization with matplotlib and seaborn.

### *Statistical Analyses*

For immunostaining, quantification statistical analyses were performed using PRISM (GraphPad, San Diego, CA). Statistical significance was calculated by one-way analysis of variance based on figure legends. The number of samples(n) analyzed were noted in figure legends and text. Data are expressed as mean ± standard deviation. For clustering analyses with Xenium data, Mann-Whitney U tests with or without Benjamini–Hochberg false discovery rate (FDR) correction were applied. Figures were generated using Adobe Photoshop and Illustrator 2026 as well as *BioRender.com*.
